# A genetic screen reveals dosage-sensitive effects of ASD genes and identifies *domino* as a regulator of synaptic and behavioral phenotypes

**DOI:** 10.64898/2026.01.12.699054

**Authors:** Ellen Stirtz, Zachary B Madaj, Brejnev Muhire, Robert Fillinger, Joe Roy, Dmitri Martirosov, Daniel Bautista, Sreeja Varanasi, Adelheid Lempradl

## Abstract

While heterozygous gene-disrupting variants in dosage-sensitive genes are strongly implicated in autism spectrum disorder (ASD), their effects on behavior *in vivo* remain poorly understood. To address this, we conducted a targeted behavioral screen in *Drosophila* using high-confidence ASD risk genes. This screen identified 48 lines with altered sleep, activity, or social behavior, including many genes not previously known to regulate these behaviors. The chromatin remodeler *domino* (*dom*) emerged as a compelling hit. Heterozygous mutants showed altered social spacing and male-biased changes in sleep and activity. RNA-sequencing revealed changes in gene expression and splicing associated with synaptic pathways. Consistent with these molecular changes, immunofluorescence revealed increased presynaptic activity in a brain region associated with sleep and sensory processing. Together, these findings show that partial loss of ASD risk genes is sufficient to alter behavior and identify *dom* as a link between transcriptional regulation, synaptic organization and behavior.

## INTRODUCTION

Autism spectrum disorder (ASD) is a multifaceted neurodevelopmental condition that typically begins early in life. According to the CDC, approximately 1 in 31 children (3.2%) are diagnosed with ASD by the age of 8(*1*). Symptoms typically emerge early in life and vary widely in severity and presentation across individuals. Phenotypic heterogeneity appears in many areas, including social interaction, communication abilities, and sleep habits(*1–3*). This phenotypic heterogeneity is mirrored at the genetic level, with hundreds of genes now implicated in ASD susceptibility(*4–6*), many of which are affected by heterozygous, gene-disrupting variants that point to dosage-sensitive mechanisms(*7*). However, for many of these risk genes, it remains unclear whether partial loss of function is sufficient to alter behavior *in vivo*. While some ASD-associated genes encode proteins directly involved in neurobiological processes such as synaptic development and neuronal signaling(*8*), others have broader regulatory roles. Among these, chromatin regulators are of particular interest, as they control the expression of large gene networks. Large-scale sequencing studies have shown that chromatin regulators are disproportionately affected by *de novo* mutations in ASD, highlighting their critical role in shaping the molecular landscape of brain development and suggesting a link between environmental signals and neuronal function(*5, 8–10*).

The growing list of candidate genes associated with ASD has created a need for scalable, functional models to investigate how molecular mechanisms influence behavior. *Drosophila melanogaster* has emerged as a powerful system for addressing this challenge(*10*). Despite evolutionary distance, flies share conserved neurodevelopmental pathways with mammals(*11–13*). Importantly, the fly model supports both high-throughput behavioral assays and precise *in vivo* genetic manipulation, making it well-suited to functionally interrogate ASD-associated genes at multiple levels.

In this study, we used *Drosophila* to screen Simons Foundation Autism Research Initiative (SFARI)(*4*) high-confidence (Category 1) ASD risk genes for their effects on sleep, activity, and social behavior (Fig. 1a). To support large-scale behavioral profiling, we refined the analysis of the locomotor activity assay(*14*) for comprehensive behavior profiling (Fig. 1b) and developed FlyFinder, an automated image analysis pipeline designed to quantify social interaction from group assays (Fig. 1c). Using a panel of *Minos*-mediated integration cassette (MiMIC)(*15*) insertion mutants targeting orthologs of SFARI Category 1 genes, we conducted our screen in heterozygous animals. This enabled us to test whether partial reduction in gene function alone is sufficient to alter behavior in our *in vivo* model, a key but largely unresolved question for ASD risk genes. While the degree of expression reduction may vary across genes and species, many high confidence ASD genes show reduced gene function in humans(*7*), making this approach appropriate for assessing their functional impact. Our screen identified 48 candidates with altered group dynamics or sleep and activity regulation in one or both sexes, including many genes not previously shown to induce behavior in any *in vivo* model.

**Fig. 1:**
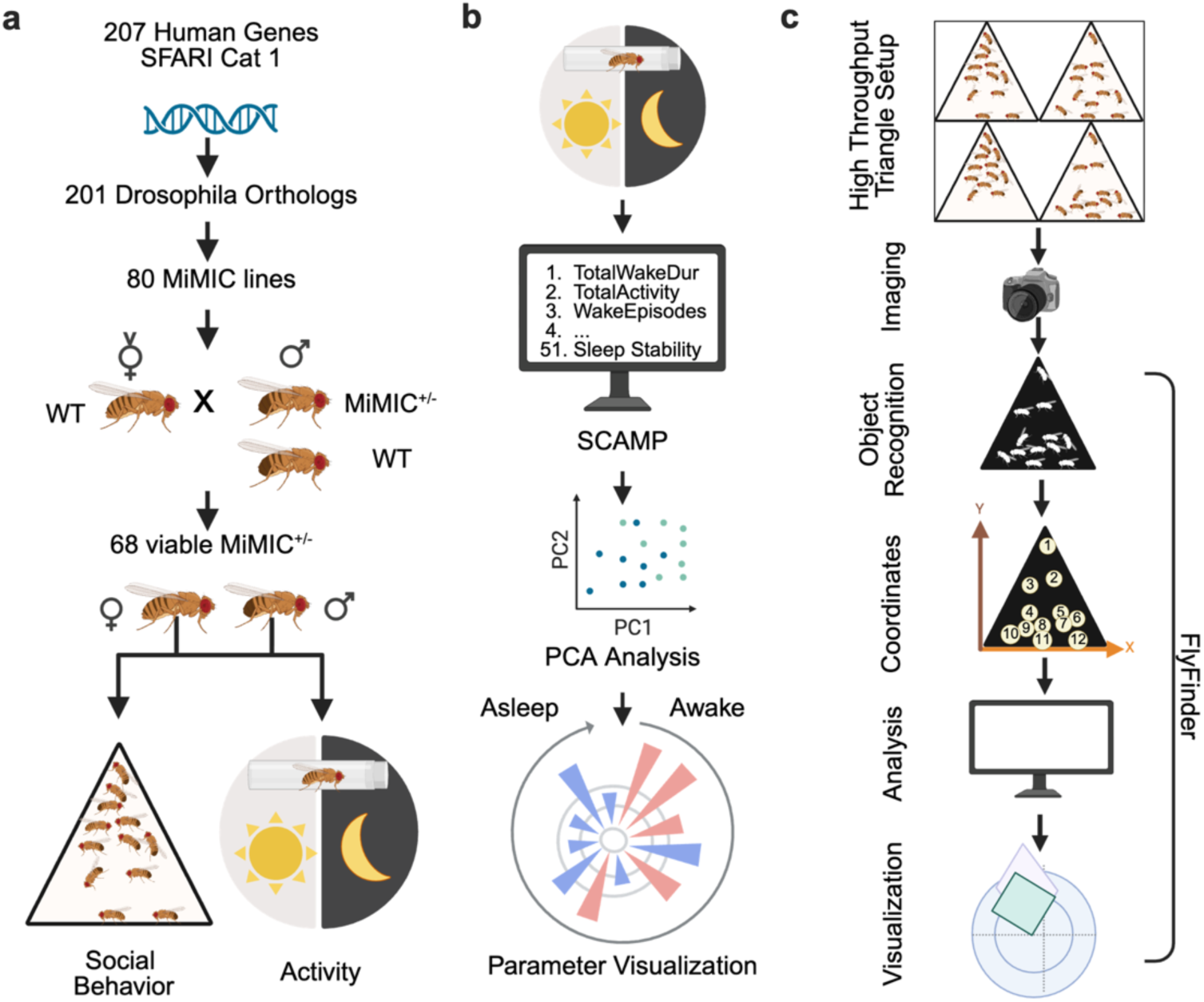
Overview of the behavioral screen and assay workflow. **a**) Schematic of the screening strategy. High-confidence ASD (Cat 1) SFARI genes were identified (207), filtered for those with a *Drosophila* ortholog (201), and their corresponding MiMIC lines were selected (80). Virgin females were crossed with wild-type or MiMIC^+/-^ males, and behavioral phenotyping was performed on male and female offspring (68 viable lines after screening). **b**) Activity assay schematic. Individual flies were monitored for 48 h using the DAM2 system, generating 51 output parameters. PCA was applied to identify global differences. Individual parameters were plotted using rose plots with parameters ordered from most wake- to most sleep-related. **c**) Social spacing assay schematic. A total of 20 flies were dispensed in a triangular arena (four arenas per trial). After 20 min, an image was captured and analyzed using FlyFinder, a custom program that automatically detects each fly, records coordinates, and calculates multiple spatial parameters (number of flies within 4 body lengths, nearest 3 neighbors, nearest neighbor, and global median).

Among these candidates, *domino* (*dom*), the fly ortholog of the human chromatin remodeler *Snf2-related CREBBP activator protein* (*SRCAP*), was found to independently modulate sleep, activity and social behavior. To understand how *dom* regulates behavior at the molecular level, we compared gene expression and RNA splicing patterns in the brains of wild-type and *dom* mutant flies. This analysis implicated synapse connectivity as an underlying structural component of the altered behavior. Indeed, immunofluorescence revealed an increase in synaptic organization in the adult *dom* mutant brains.

Together, our screen identifies regulators of sleep and social behavior. By assessing heterozygous loss, we show that partial reduction of ASD risk genes is sufficient to alter behavior *in vivo* and link the chromatin remodeler Domino to synaptic changes.

## RESULTS

### Identification of conserved candidate ASD genes in *Drosophila*

To identify ASD-relevant genes that affect behavior, we performed a targeted genetic screen in *Drosophila*, focusing on sleep, activity and social interaction, core characteristics often affected in autistic individuals (Fig. 1a). At the time of study initiation, the SFARI GENE database listed 207 Category 1 human genes, each with strong evidence for association with ASD, typically based on the presence of at least three *de novo* likely gene-disrupting mutations(*4*). Of these, 201 had identifiable *Drosophila* orthologs (DRSC integrative ortholog prediction tool)(*16*). To ensure a consistent genetic background, we utilized the MiMIC gene disruption collection, which consists of transposon-based insertions into intronic regions that disrupt gene function through premature termination(*15*). The mutagenic cassette contains a splice acceptor followed by stop codons in all three reading frames. Because the mutagenic splice acceptor and stop cassette must be oriented in the same transcriptional direction as the host gene to effectively disrupt gene function, we selected lines with correctly oriented insertions. This yielded 80 strains targeting 97 high confidence SFARI genes (Ext. Table 1), with some fly orthologs mapping to multiple human SFARI genes.

**Table 1.** Genetic screen hits.

| Sleep and Activity |  |  | Social Behavior |  |  | Overlap |  |
| --- | --- | --- | --- | --- | --- | --- | --- |
| Males | Females | Both | Males | Females | Both | Males | Females |
| *bab1* | *Ank2* | *bchs* | *bel* | *Ank2* | *bel* | *dom* | *Ank2* |
| *bchs* | *bchs* | *brm* | *CG43367* | *bel* | *Sin3A* | *Nlg3* | *brm* |
| *brm* | *brm* | *Ekar* | *cic* | *brm* |  | *Nrx-1* | *Sin3A* |
| *bru3* | *CG17819* | *KCNQ* | *dom* | *hbn* |  | *Prosap* |  |
| *btv* | *CG43367* | *Nlg3* | *kn* | *Hr3* |  |  |  |
| *CG10984* | *Doc1* | *Prosap* | *Lpt* | *Nhe2* |  |  |  |
| *DIP2* | *Ekar* |  | *Nlg3* | *Sin3A* |  |  |  |
| *dom* | *fne* |  | *Nrx-1* |  |  |  |  |
| *Ekar* | *FoxP* |  | *osa* |  |  |  |  |
| *Fife* | *KCNQ* |  | *Prosap* |  |  |  |  |
| *fus* | *Kdm2* |  | *Sin3A* |  |  |  |  |
| *GluRIB* | *kis* |  |  |  |  |  |  |
| *hbn* | *kn* |  |  |  |  |  |  |
| *Hr3* | *Nlg3* |  |  |  |  |  |  |
| *KCNQ* | *Prosap* |  |  |  |  |  |  |
| *LBR* | *Sin3A* |  |  |  |  |  |  |
| *Nhe2* | *Syn* |  |  |  |  |  |  |
| *Nlg3* | *tai* |  |  |  |  |  |  |
| *Nrx-1* | *trio* |  |  |  |  |  |  |
| *otk* | *wrd* |  |  |  |  |  |  |
| *pan* |  |  |  |  |  |  |  |
| *Prosap* |  |  |  |  |  |  |  |
| *Rdl* |  |  |  |  |  |  |  |
| *rn* |  |  |  |  |  |  |  |
| *Shab* |  |  |  |  |  |  |  |
| *siz* |  |  |  |  |  |  |  |
| *skd* |  |  |  |  |  |  |  |
| *Tie* |  |  |  |  |  |  |  |
| *Ube3a* |  |  |  |  |  |  |  |
| *Vps13B* |  |  |  |  |  |  |  |

Among the genes analyzed, the largest functional groups were those involved in neurodevelopment and synaptic function (19) and transcriptional regulation (19), followed by genes associated with cell organization and cytoskeletal dynamics (13), epigenetic regulation and chromatin remodeling (11), transport and localization (9), and signaling or receptor activity (6). An additional 3 genes did not fall into a specific functional category. A total of 12 lines yielded insufficient viable offspring and were excluded from the final analysis. We generated heterozygous animals for phenotyping by crossing male MiMIC mutants with wild-type females (Fig. 1a). Since MiMIC insertions generally produce strong loss of function alleles, the resulting progeny represent heterozygous loss of function mutants, modeling partial gene disruption. This design allows us to test whether reduced gene dosage alone is sufficient to alter behavior *in vivo*. In addition, this approach minimizes confounding background effects and enables a systematic comparison across a large set of ASD risk genes in a highly controlled manner.

### Genetic screening in *Drosophila* identifies known and novel modulators of sleep and activity

To explore changes in sleep and activity, we measured movements of 3 to 5-day-old male and female flies over a 48 h period using the DAM2 Drosophila Activity Monitor(*14*). Data were analyzed using SCAMP(*17*), which calculates 51 different measures of sleep and activity behavior (Fig. 1b). To identify patterns across behavioral variables that distinguish wild-type and mutant flies, we performed principal component analysis (PCA) on all 51 measurements. This analysis identified a subset of genes with robust, mutant-specific effects on sleep and activity.

PCA was conducted separately for males and females for each mutant line and its paired control. Linear mixed-effects models were used to test how each of the first 6 principal components (PCs), which together capture ≥80% of the variance in both sexes (Ext. Table 2), differed across genotypes. Breeding vial was included as a random effect to account for similarity among flies in shared rearing conditions (3-4 flies per vial per testing period). This analysis revealed consistent patterns within PCs across genotypes (heatmap in Fig. 2a,b; left panel), indicating systematic effects of genetic background, particularly for male comparisons. To identify mutant-specific behavioral signatures that were independent of these MiMIC-associated background effects, we estimated 95% confidence intervals and calculated second-generation p-values (SGPVs) using a ±10% null interval for two complementary comparisons: (i) each mutant genotype vs wild-type; and (ii) each genotype vs the pooled mean of all other mutant lines. Genes meeting both criteria (SGPV = 0 for both) were classified as hits (red), whereas genotypes differing from wild-type only were classified as background-associated (gray) (Fig. 2a,b; right panels). In males, PCs 1, 2, and 4, and in females, PCs 1-4, were dominated by wild-type only effects, evident as consistent shading across genotypes in the heatmaps and a higher proportion of gray boxes (Fig. 2a,b). The same pattern was also apparent in the corresponding forest plots (Supplementary Fig. 1a,b), which rank genes within each PC by their estimated effect size. In these plots, genes classified as hits (red) were enriched at the extremes of the distributions, whereas wild-type only effects (gray) were distributed more broadly across effect sizes. In contrast, the remaining PCs, PCs 3, 5, and 6 in males, and PCs 5 and 6 in females, contained relatively few genotypes with wild-type only effects, suggesting that variation along these axes is driven primarily by mutant-specific effects rather than shared background influences.

**Fig. 2:**
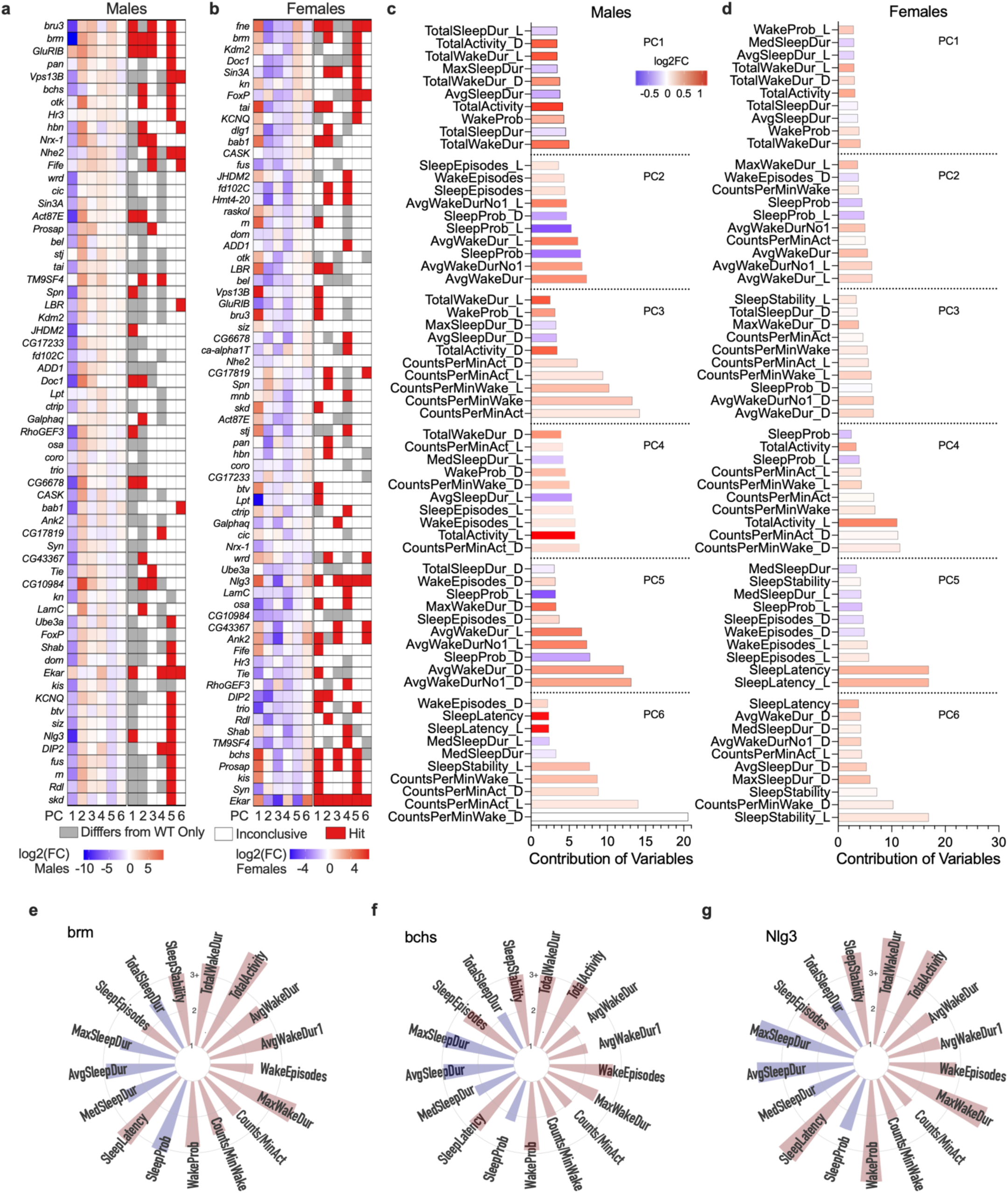
Genetic screening in Drosophila identifies modulators of sleep and activity. **a,b**) Log2fold change heatmaps and significance plots for all 68 viable screened genes in males (**a**) and females (**b**). For the top 6 PCs, 95% confidence intervals were computed for two comparisons: (i) each mutant genotype versus wild-type and (ii) each genotype versus the pooled mean of all other mutant lines. SGPVs were calculated for each comparison. Genes that met hit criteria (SGPV = 0) for the wild-type comparison alone are shown in gray, whereas genes that met both criteria are shown in red. Rows are ordered by the estimated effect size for PC5 and displayed in descending order (n=2-3 replicates per line with 4 flies per replicate). **c,d**) Bar graphs show the relative contribution of each behavioral parameter to the indicated PC, expressed as a percentage of the total contribution for that component, shown separately for males (**c**) and females (**d**). Color indicates the mean log2fold change between mutants and wild-type, with red denoting higher values in mutants and blue denoting higher values in wild-type. Y-axis labels denote individual behavioral parameters, with “D” indicating dark-phase measurements and “L” indicating light-phase measurements. **e-g**) Rose plots of top genes significantly different by both criteria and shared between sexes, shown for males. Plots are organized clockwise from “more awake” to“more asleep” and summarized across the full 24 h monitoring period. Red bars indicate increased behavior in MiMIC^+/-^ flies, while blue bars indicate decreased behavior.

Focusing on variable distribution within individual PCs, those dominated by wild-type only effects exhibited diffuse loadings, reflected by a lack of a clear structure in the corresponding contribution plots (Fig. 2c,d). In these plots, bar height reflects the relative contribution of each variable to the PC, while color indicates the direction and magnitude of the pooled mutant vs wild-type log2fold change for that variable. By contrast, PCs associated with mutant-driven variation showed much more specific and interpretable variable loadings. In males, PC3 reflected a general increase in activity during wakefulness across the full 24 h period, while PCs 5 and 6 captured phase-specific waking changes driven predominantly by the dark (7pm–7am) or light (7am–7pm) phases, respectively (Fig. 2c). In females, PC5 was largely driven by light-phase sleep latency, or the time required to return to sleep after waking, while PC6 captured increased waking activity during the dark phase alongside changes in sleep stability (Fig. 2d). These distinctions illustrate how individual mutant-driven PCs capture different combinations of behavioral features and temporal dynamics.

Based on these results, we designated hits along PCs 3, 5, and 6 in males and PCs 5 and 6 in females as our primary readout of genetic effects. Using this criterion, we identified 30 significant hits in males and 20 in females (Ext. Table 2), including both known and novel regulators of sleep and activity across the full range of functional categories represented in the screen (Ext. Table 1). Across sexes, 6 genes emerged as common hits (*bchs, brm, Ekar*, KCNQ, *Nlg3*, and *Prosap*), highlighting a core set of genes with conserved behavioral effects. To provide a consolidated view of behavioral outcomes, we summarized results across assays in Table 1. Of note, even within background-dominated PCs, mutants differed from controls in distinct ways. Some mutants exhibited exaggerated versions of the background behavioral pattern (e.g., *brm* on PC1 in males; a dark blue box in Fig. 2a and a mean difference at the negative extreme of Supplementary Fig. 1a), whereas others reversed the background trend entirely (e.g., *GluRIB* on PC1 in males; a light red box in Fig. 2a and a mean difference at the positive extreme of Supplementary Fig. 1a). Together, these patterns illustrate multiple modes by which genetic perturbations in ASD-associated genes can shape sleep- and activity-related behavior in *Drosophila*.

To connect these PC-level signatures to individual genes, we examined representative hits in greater detail. An example of a known regulator of the circadian clock is *brahma (brm),* which has been linked to increased wakefulness when disrupted(*18*). Consistent with this role, *brm* was classified as a hit in PC3 and PC5 in males, two components associated with wake-related variables (Fig. 2a,c). Rose plots visualizing all behavioral parameters across the 24 h recording period provide a more granular view of the changes underlying these PC-level effects. In males, these plots highlighted increased wake duration, elevated overall activity, and prolonged sleep latency relative to wild-type controls (Fig. 2e). Similarly, *brm^+/-^* females showed heightened sleep latency (Supplementary Fig. 1c), consistent with its classification as a hit in PC5. As a second example, *blue cheese (bchs),* which has previously been reported to affect sleep(*19*), was classified as a hit in PC5 in both sexes and shows altered patterns of wakefulness throughout the monitoring period (Fig. 2a,b,f and Supplementary Fig. 1d). Beyond validating known regulators, the screen also highlighted novel candidates. *Neuroligin 3* (*Nlg3),* for example, was classified as a hit in PC5 in males and PCs 5 and 6 in females, which are driven by activity measures and sleep-related variables, respectively. Rose plots for *Nlg3^+/-^* highlight alterations in wake duration and total activity in both males and females (Fig. 2g and Supplementary Fig. 1e). Although *Nlg3* itself has not been directly linked to sleep regulation, mutations in the related gene *Neuroligin 4* (*Nlg4)* have been shown to reduce sleep and increase nighttime awakenings(*20*). Our results therefore suggest that *Nlg3* influences behavioral state regulation in a manner that might be conserved across neuroligin family members and manifest differently between males and females. Overall, these results demonstrate that the screen both validates known regulators of sleep and identifies novel candidates, supporting its utility for uncovering the role of ASD-relevant genes in modulating global patterns of sleep and activity.

### Genetic screening in *Drosophila* identifies modulators of social behavior

To assess social behavior in a group setting, we utilized an established social spacing assay(*21*) for scalable, automated analysis (Fig. 1c). The assay quantifies how individual flies position themselves relative to others, providing a measure of social organization. To adapt the assay for high throughput experiments, we first expanded the setup to include four triangles, allowing parallel testing of multiple groups (e.g., MiMIC^+/-^ vs wild-type or male vs female) and thereby increasing throughput and experimental consistency. We then developed a custom image segmentation program, FlyFinder (https://github.com/LempradlLab/drosophila_screen_dom), which automatically separates the four-triangle images and extracts individual fly coordinates to calculate multiple spatial parameters. This automated approach eliminates manual bias and enables high-throughput, reproducible quantification of social interactions.

Our analysis included a standard measure of social spacing as described previously(*22*), the number of flies within 4 body lengths, which captures interactions within a small group of individuals. We also measured the nearest neighbor distance, which reflects the proximity to a single fly; the median distance to the 3 nearest neighbors, which helps distinguish loose groups from tight clusters; and the global median distance, which can be interpreted as an indicator of overall group distribution. Together, these parameters provide complementary perspectives on group cohesion and social preference. Each mutant line was measured in parallel with its corresponding wild-type control, ensuring that results were not confounded by external testing conditions. For each outcome, we calculated the same two SGPVs used in the activity analysis, (i) comparing the genotype with wild-type and (ii) comparing between all other mutant lines. Genes were considered hits if they exhibited significant SGPVs (SGPV = 0) for both comparisons.

Using this approach, we identified 11 genes in males and 7 genes in females that play a significant role in regulating social behavior (Fig. 3a,b; red boxes, right panels), further underscoring the behavioral relevance of many ASD-associated genes. These hits collectively converge on epigenetic, transcriptional, and synaptic functions. Overlap between sexes was limited, with only 2 genes (*bel* and *Sin3A)* emerging as top hits in both males and females (Table 1 and Ext. Table 2). Most hits were associated with tighter clustering of flies (observed in 2/3 male hits and all female hits), evident as red shading across genotypes for the number of flies within 4 body lengths and blue shading for the remaining distance-based parameters (Fig. 3a,b; heatmaps, left panels). This shift in group organization is further illustrated in the forest plots (Supplementary Fig. 2a,b), where most hits reside to one extreme of the distribution, consistent with increased social proximity, suggesting that many of the identified genes converge on shared features of social organization and produce similar shifts in group structure across sexes.

**Figure 3:**
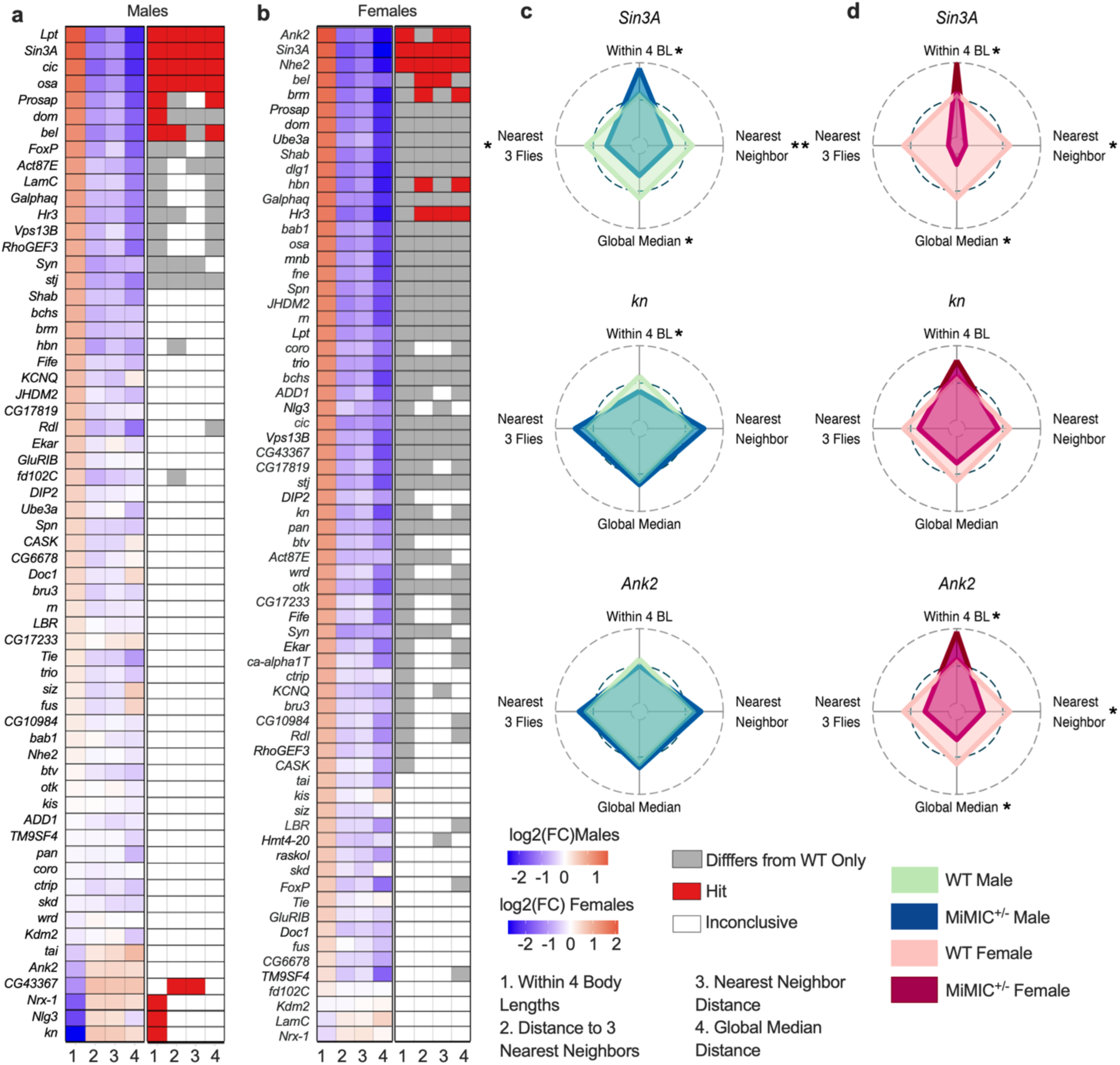
Genetic screening in Drosophila identifies modulators of social behavior. **a,b**) Log2fold change heatmaps and significance plots for all screened genes are shown for the four social spacing parameters in males (**a**) and females (**b**). SGPVs were calculated for the comparisons (i) genotype vs wild-type and (ii) genotype vs all other mutant lines. Genes that met hit criteria (SGPV = 0) for the wild-type comparison alone are shown in gray, whereas genes that met both criteria are shown in red. Rows are ordered by the estimated effect size for Within 4 Body Lengths and displayed in descending order (n=2-3 replicates per line with 4 flies per replicate). **c,d**) Radar plots of the top shared (top), male-specific (middle), and female-specific (bottom) significant genes, showing t-statistics from the mutant vs wild-type contrasts in males (**c**) and females (**d**). The inner polygon represents the wild-type reference (t = 0), with axes extending outward indicating higher values relative to wild-type and axes plotting inward indicating lower values. Male and female plots share a common axis scale to allow direct comparison across genotypes.

Radar plots (Fig. 3c,d) provide a more detailed view of individual lines, with darker shaded regions representing the mutant behavior profiles. *SIN3 transcription regulator family member A (Sin3A)* was the strongest shared hit across sexes, showing significantly tighter clustering in both males and females (Fig. 3c,d; top panels). As expected, local clustering (number of flies within 4 body lengths) was inversely correlated with nearest neighbor measurements, such that tighter clustering was associated with shorter distances to the closest neighbor. Likewise, the three nearest neighbor and global median distance metrics suggest that wild-type flies tend toward a more disperse group organization. Notably, *Sin3A* encodes a chromatin regulator implicated in Witteveen-Kolk syndrome (OMIM 613406), a neurodevelopmental disorder often associated with ASD(*23*), underscoring the utility of this *Drosophila* model for identifying functionally relevant ASD risk genes. Among sex-specific hits, *knot (kn)*, the ortholog of human *EBF3*, a transcriptional regulator of nervous system development(*24, 25*), was identified as a male-specific hit associated with significantly increased social spacing (Fig. 3c; middle panel). In females, *Ankyrin 2 (Ank2),* which is important for axonal function and synaptic connectivity(*26*), drove significantly decreased social spacing (Fig. 3d; bottom panel). Taken together, these findings demonstrate that ASD-associated genes spanning diverse molecular functions play important roles in shaping group organization, pointing to a subset of high-confidence candidates warranting further experimental validation.

### Partial loss of *domino* alters social spacing, sleep, and activity in a sex-specific manner

Among the genes identified in the screen, the chromatin remodeler *domino* (*dom*) stood out as one of the few candidates affecting both behavioral domains. Specifically, flies heterozygous for the MiMIC insertion within *dom*(*27*) (Fig. 4a; dom^MiMIC+/-^ flies) exhibited male-specific alterations in sleep, activity, and social spacing (Fig. 2a,b and Fig. 3a,b). *dom* is the *Drosophila* ortholog of human *Snf2-related CREBBP activator protein* (*SRCAP*), a core component of a chromatin remodeling complex that incorporates H2A.Z-H2B dimers into nucleosomes to facilitate transcription(*28*). Together with the established role of chromatin remodeling in neural development and its genetic linkage to ASD(*29, 30*), *dom* represents a strong candidate.

**Fig. 4:**
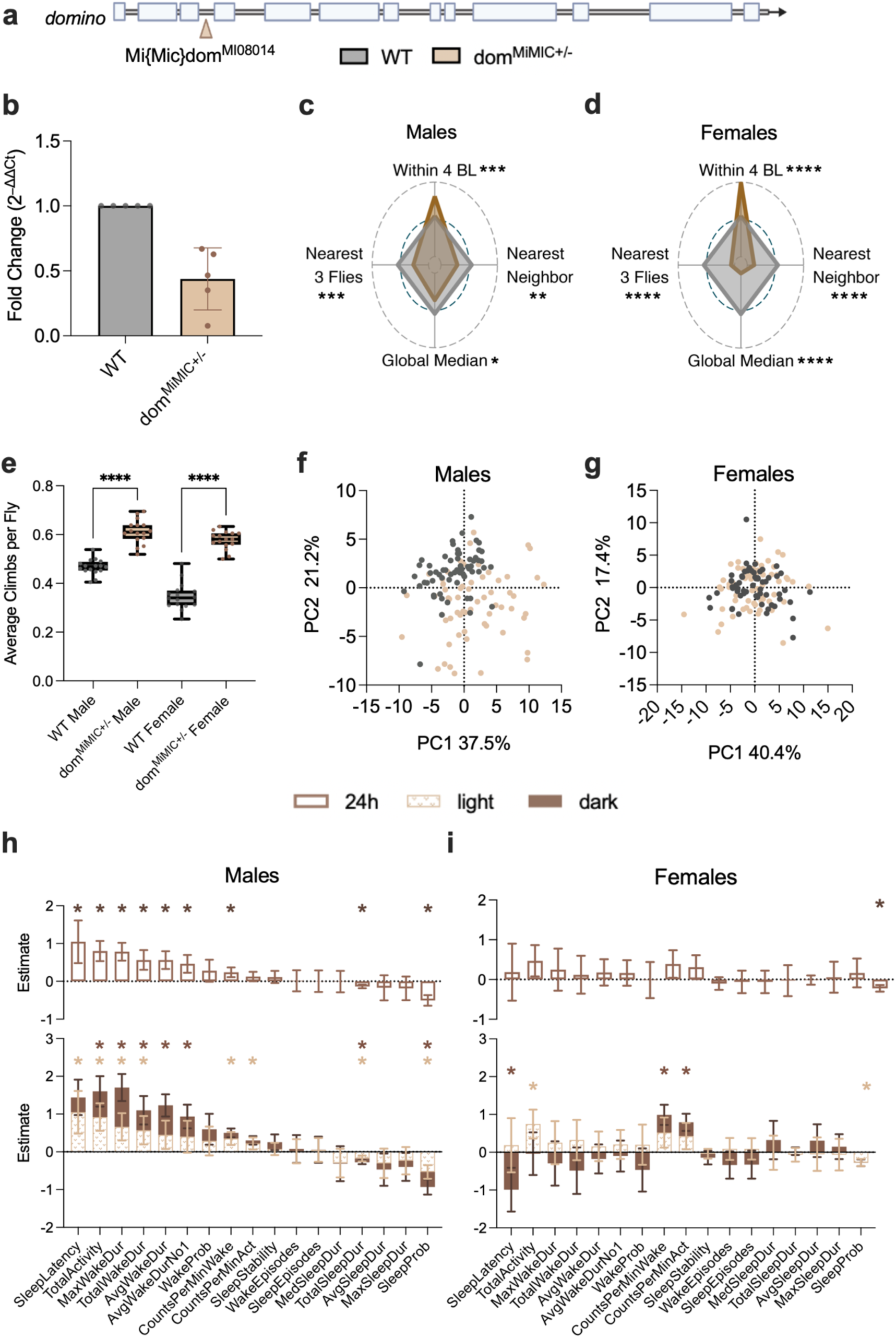
Behavioral characterization of dom^MiMIC+/-^ flies. **a**) Schematic of the domino^MIMIC+/-^ insertion. **b**) qPCR analysis of dom expression in wild-type and dom^MiMIC+/-^ flies (n=5; 10 heads per replicate; Kruskal-Wallis test). **c,d**) Radar plots of social spacing assay results for males (**c**) and females (**d**), showing t-statistics from the *dom* vs wild-type contrasts (n=16; 20 flies per replicate). The inner polygon represents the wild-type reference (t = 0), with axes extending outward indicating higher values relative to wild-type and axes plotting inward indicating lower values. Male and female plots share a common axis scale to allow for direct comparison. **e**) Climbing assay performance in males and females (n=16; Two-way ANOVA with Šídák multiple comparisons). **f,g**) PCA of activity assay results in males (**f**) and females (**g**), with wild-type (gray) and dom^MiMIC+/-^ (tan) flies (n=64). **h,i**) Individual sleep and activity parameters for males (**h**) and females (**i**). For each measure, the 24 h estimate is shown in the upper graph (open bars), with light- and dark-period estimates shown in the lower graph (patterned and filled bars, respectively). Estimates are plotted such that positive values indicate an increase and negative values indicate a decrease in dom^MiMIC+/-^ flies relative to controls. Asterisks indicate significance (p ≤ 0.01), with light and dark brown asterisks denoting significance during the light and dark periods, respectively.

To confirm that the MiMIC allele results in the disruption of gene function, we assessed transcript levels in heterozygous flies. qPCR analysis of heads shows a ∼50% reduction in expression (Fig. 4b), consistent with partial loss of gene dosage. We next repeated the phenotyping experiments with additional replicates to increase statistical power relative to the initial screen. Under these conditions, both male and female dom^MiMIC+/–^ flies exhibited behavioral effects in the social spacing assay, consistent with tighter clustering (Fig. 4c,d and Ext. Table 3). To rule out the possibility that this phenotype was a secondary consequence of impaired locomotor ability, given that the assay requires vertical climbing, we performed Rapid Iterative Negative Geotaxis (RING) tests. These showed no motor impairment; in fact, climbing ability was increased in both male and female mutants (Fig. 4e).

We next sought to confirm the sleep and activity patterns observed in the primary screen. Principal component analysis showed that PC1 and PC2 clearly separated dom^MiMIC+/-^ males from wild-type controls, accounting for ∼38% and ∼21% of the variance, respectively (Fig. 4f), whereas female dom^MiMIC+/-^flies showed no separation along the first two PCs (Fig. 4g). This confirms the male-biased activity differences suggested in the primary screen. We next examined individual sleep and activity measures across the 24 h monitoring interval (Fig. 4h,i; top panels) and separated by light and dark periods (Fig. 4h,I; bottom panels). Male dom^MiMIC+/-^ flies exhibited significant changes in sleep-related parameters, including increased latency to fall asleep, reduced sleep duration, prolonged wakefulness and decreased sleep probability. While changes in sleep were observed during both light and dark periods, increased activity was restricted to daytime (Fig. 4h). These parameters have established relevance to ASD-associated clinical features, including sleep latency, efficiency, fragmentation, and hyperactivity(*31*). Female dom^MiMIC+/-^ flies, by contrast, showed fewer overall differences, with significant effects primarily restricted to activity-related parameters (Fig. 4i). Together, these findings indicate sex-dependent effects of *dom* heterozygosity, with broad changes in sleep and activity-related parameters in males and limited activity-related effects in females.

To validate the *dom*-dependent effects observed in the original fly line, we analyzed a second allele, dom^Trojan+/-^, in which a Mi{Trojan-GAL4.0} cassette is inserted into the same genomic site as the dom^MiMIC+/-^line(*32*) (Supplementary Fig. 3a). This insertion causes an even more pronounced (∼75%) reduction in expression in heterozygous animals (Supplementary Fig. 3b). Behavioral analysis largely mirrored the dom^MiMIC+/-^ phenotypes (Ext. Table 3): dom^Trojan+/-^ flies clustered more tightly in the social spacing assay (Supplementary Fig. 3c,d), exhibited enhanced climbing ability (Supplementary Fig. 3e), and largely mirrored activity and sleep patterns of dom^MiMIC+/-^ flies (Supplementary Fig. 3f-i). Notably, in the activity assay, dom^Trojan+/-^ females showed slightly better separation from controls by PCA and displayed more pronounced changes across individual sleep and activity parameters (Supplementary Fig. 3g,i). Together with the MiMIC data, these results demonstrate that effects were reproduce across independent alleles, supporting a robust role for *dom* in regulating behavior.

To assess the cellular origin of these phenotypes, we next tested whether cell-specific depletion of *dom* could reproduce the observed phenotypes. We performed RNAi knockdown using GAL4 drivers targeting the whole body (tubulin-Gal4), all neurons (elav-Gal4), mushroom body neurons (MB-Gal4) and glial cells (repo-Gal4). At the pupal stage, only whole-body knockdown resulted in lethality, with the other crosses showing no significant differences (Supplementary Fig. 3j-m). When examining adult survival, no flies eclosed with whole-body, pan-neuronal, or glial knockdown (Supplementary Fig. 3n-p). Mushroom body knockdown allowed limited survival to adulthood, but these rare survivors were severely impaired: they could not climb or fly and died shortly after eclosion (Supplementary Fig. 3q), precluding detailed phenotyping. These results are consistent with previous reports of *dom* lethality outside of embryogenesis(*33, 34*) and indicate that substantial depletion of *dom*, both globally and in specific brain cells, limits adult viability, confirming that heterozygosity is appropriate to assess the behavioral impact of *dom* loss.

### *Domino* heterozygosity alters transcriptional programs involved in synaptic signaling

Given that *dom* encodes a chromatin remodeler, we next assessed gene expression using RNA sequencing (RNA-seq) on 3 to 5-day-old adult heads, corresponding to the stage at which behavioral phenotypes were assessed. Despite its role as a transcriptional activator(*33, 35*), we did not observe a consistent bias toward downregulation. Differential expression analysis identified 244 significantly altered genes in dom^MiMIC+/-^ males (Fig. 5a and Ext. Table 4; 140 down-, 104 upregulated) and 560 in dom^MiMIC+/-^ females (Fig. 5b and Ext. Table 4; 147 down-, 413 upregulated). As expected, *dom* expression itself was significantly reduced in both sexes, with log2 fold change values closely matching qPCR measurements (Fig. 5a,b), thereby confirming the heterozygous loss-of-function mutation.

**Fig. 5:**
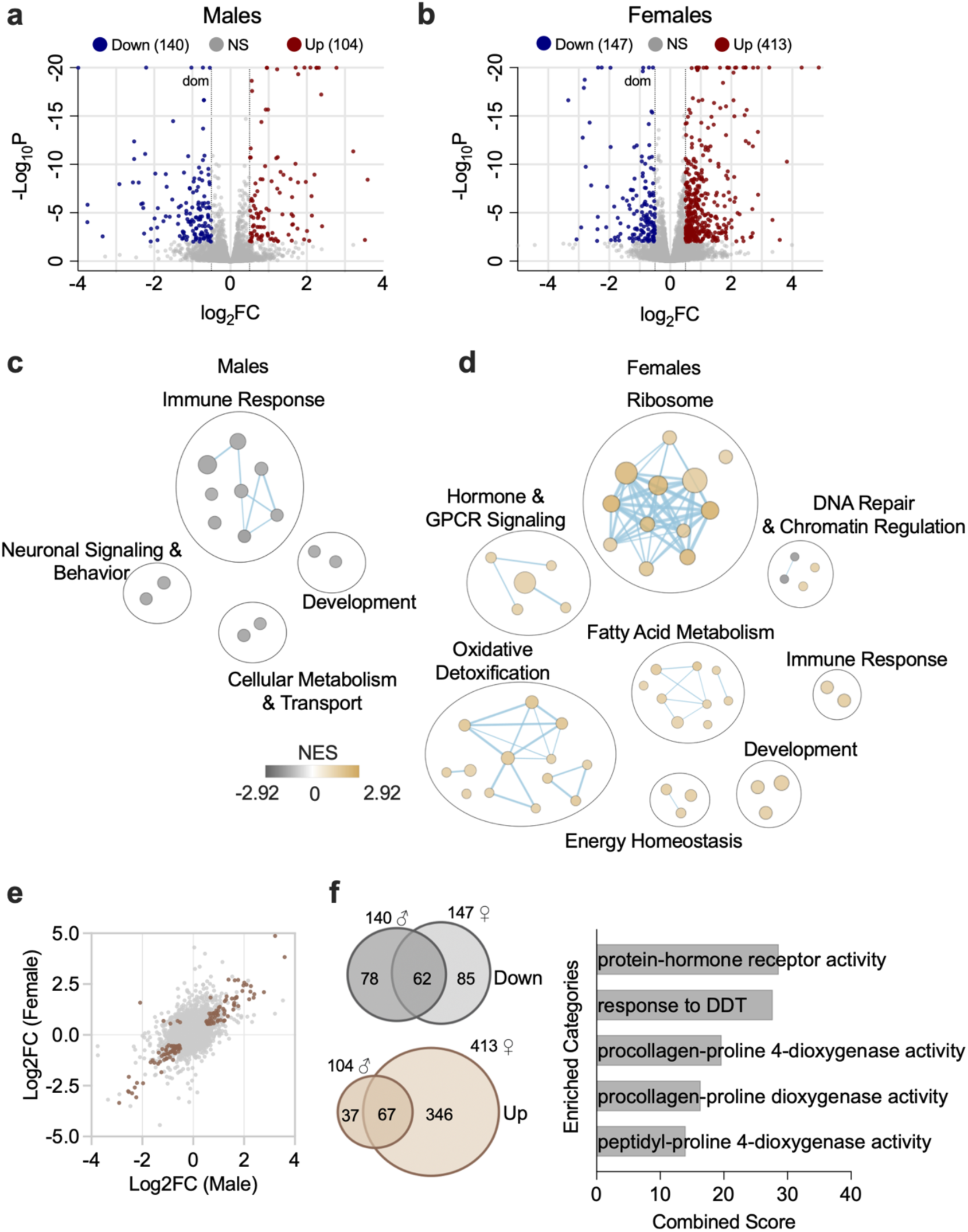
domMiMIC^+/-^ flies exhibit distinct transcriptional profiles in males and females. **a,b**) Volcano plots of DE genes in males (**a**; 140 downregulated and 104 upregulated), and females (**b**; 147 downregulated and 413 upregulated) (n=5 replicates of 5 pooled brains; padj ≤ 0.01 and |log2fold change| ≥ 0.5). Expression of *dom* is significantly reduced in both sexes. **c,d**) GSEA of all DE genes in males (**c**) and females (**d**). **e**) Scatterplot of overlapping significant DE genes in males and females (tan), plotted by log2fold change (overlap significance, Fisher’s Exact Test <0.0001). **f**) Venn diagrams of downregulated (gray) and upregulated (tan) genes in males (left) and females (right). Numbers above circles indicate the total number of DE genes per condition, while numbers within circles denote unique or shared genes across sexes. ORA of shared genes is shown in the adjacent bar plots (padj ≤ 0.05).

To identify pathway level patterns in the RNA-seq dataset, we applied Gene Set Enrichment Analysis (GSEA)(*36, 37*) to normalized expression data, allowing the detection of coordinated changes across predefined sets of genes. The resulting GSEA terms were visualized in Cytoscape(*38*), where related pathways cluster by functional similarity (Fig. 5c,d; a complete list of pathways and enrichment scores is available in Ext. Table 4). In dom^MiMIC+/-^ males, genes associated with neurotransmitter transport and olfactory behavior were downregulated (Ext. Table 4), suggesting reduced neuronal signaling capacity (Fig. 5c). In females, enrichment was observed for hormone- and cell surface receptor signaling pathways (Ext. Table 4), indicating changes in intercellular and hormonal communication (Fig. 5d). Although the specific categories differed between sexes, both analyses implicated processes that influence synaptic communication. Males and females also shared a significant number of dysregulated genes, with expression changes largely correlated in direction (Fig. 5e). Across both sexes, 62 genes were commonly downregulated and 67 upregulated (Fig. 5f). Overrepresentation analysis (ORA) of these overlapping genes using FlyEnrichr(*39, 40*) highlighted pathways involved in hormone receptor activity and procollagen formation (Fig. 5f and Ext. Table 4). Hormone receptor activity reflects altered neuromodulatory signaling, while procollagen formation contributes to extracellular matrix assembly, a process critical for neuronal migration, axon guidance, and synapse formation(*41–43*). Together, these results indicate that shared transcriptional changes across sexes converge on pathways regulating synaptic signaling and structure, offering a molecular basis for the behavioral phenotypes observed in *dom* heterozygotes.

The additional dom^Trojan+/-^ fly line showed highly similar transcriptomic profiles (Supplementary Fig. 4a-f), with both sexes showing enrichment of numerous hormone- and receptor-mediated signaling pathways (Supplementary Fig. 4c,d and Ext. Table 4). These findings demonstrate that the two *dom* mutant lines produce highly similar transcriptomic signatures, reinforcing the conclusion that partial loss of *dom* disrupts signaling-related processes across sexes and highlights synaptic communication as a shared point of vulnerability.

### *Domino* heterozygosity leads to altered splicing patterns in the brain

Beyond its role in gene expression regulation, histone variant H2A.Z has been shown to maintain splicing integrity in yeast, and its loss is associated with differential splicing and disruption of neuronal and synaptic signaling in memory-related contexts in mice(*44, 45*). To determine whether partial loss of *dom* similarly affects RNA processing in *Drosophila*, we first examined splicing in our RNA-seq dataset from adult brains.

Analysis using rMATs(*46*), a tool that detects differential alternative splicing from RNA-seq datasets, revealed widespread alterations across all five major splicing event types in the brains of male and female dom^MiMIC+/-^ flies (Fig. 6a,b, Ext. Table 5). ORA of genes with significant splicing changes showed strong enrichment for pathways involved in synaptic signaling, neuronal maturation, and development (Fig. 6c,d). Consistent with these findings, dom^Trojan+/-^ flies exhibited highly similar splicing alterations (Supplementary Fig. 5a-d, Ext. Table 5), indicating that these effects are robust across independent alleles.

**Fig. 6:**
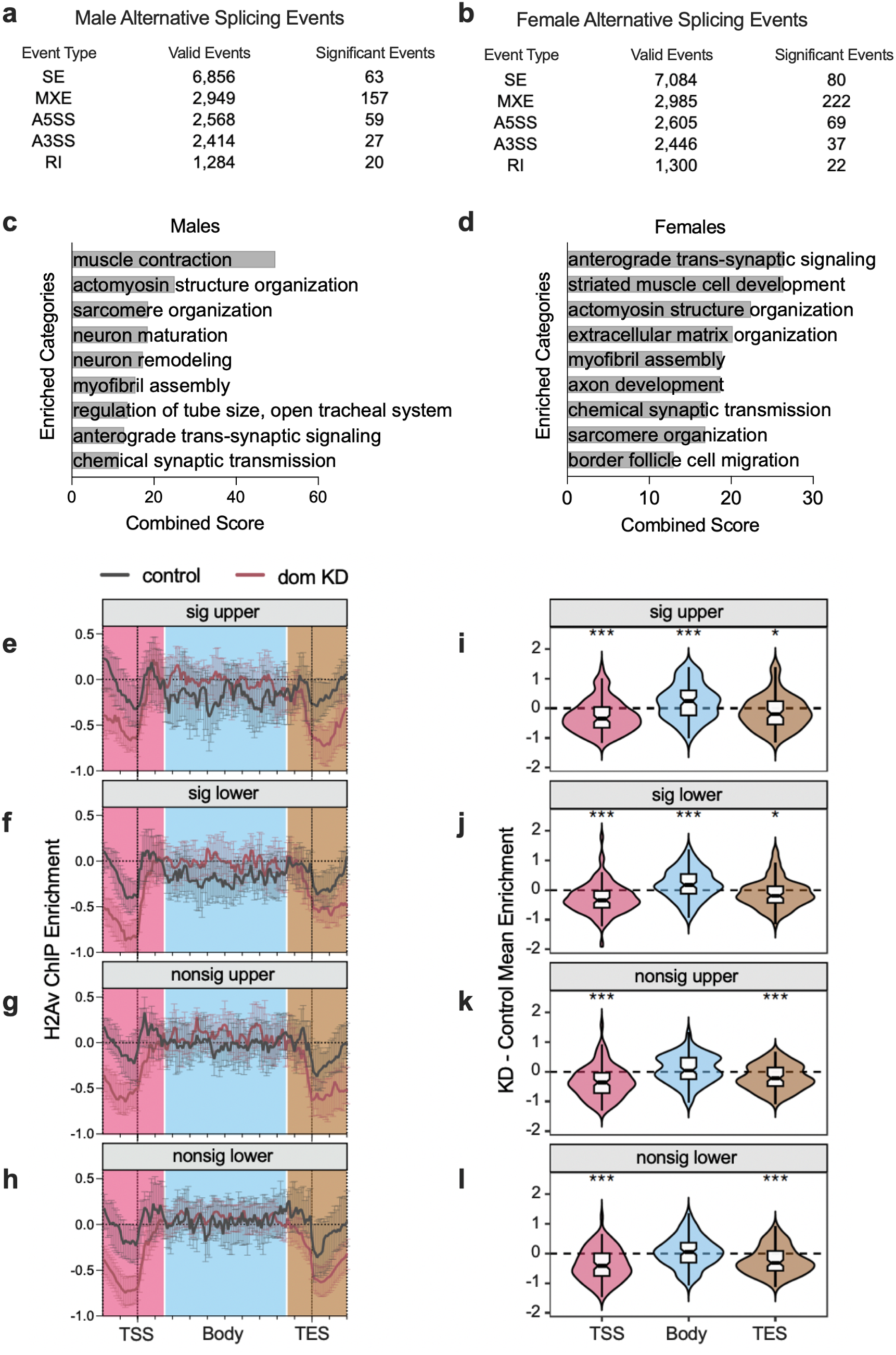
Partial loss of domino alters splicing regulation and H2A.V occupancy. **a,b**) Alternative splicing events detected in male (**a**) and female (**b**) adult dom^MiMIC+/-^ brains (n=5 replicates of 5 pooled brains), showing total valid events detected by rMATS and significant events (FDR ≤ 0.05 and |IncLevelDifference| ≥ 0.1). **c,d**) ORA of significant splicing events in males (**c**) and females (**d**). **e-g**) H2A.V ChIP enrichment profiles for control (gray) and *dom* knockdown (pink) embryos. Plots show genes with significant differential splicing identified by rMATs(*46*), divided into equally sized groups representing the upper (**e**) and lower (**f**) halves of the ranked gene set. Two matched groups of non-significant genes, ranked immediately below the splicing significance threshold, are shown for comparison (**g**), top half and (**h**), bottom half. Pink shading marks TSS, blue shading indicates gene bodies, and tan shading marks TES. **i-l**) Quantification of the mean difference in H2A.V enrichment between control and *dom* knockdown embryos (Two-way ANOVA mixed-effects analysis).

To assess whether H2A.V (H2A.Z ortholog) occupancy is associated with splicing outcomes, we reanalyzed a published dataset(*47*), combining H2A.V chromatin immunoprecipitation sequencing (ChIP-seq) and RNA-seq from zygotic genome activation (ZGA)-stage embryos following *dom* knockdown. Using rMATs, we identified widespread splicing differences between wild-type and *dom* knockdown embryos (Ext. Table 6). To assess whether H2A.V occupancy differs between genes affected by splicing changes and those that were not, we divided genes into four equally sized groups: the upper and lower halves of significantly spliced genes and two matched groups of non-significant genes ranked immediately below the splicing significance threshold. ChIP-seq profiles showed that *dom* knockdown broadly reduced H2A.V occupancy at transcription start sites (TSS) and transcription end sites (TES) across all four gene groups (Fig. 6e-l). In contrast, differently spliced genes exhibited increased H2A.V enrichment across their gene bodies (Fig. 6e,f,i,j; blue region). This pattern suggests a redistribution of H2A.V from TSS and TES proximal regions into gene bodies following *dom* disruption. Although derived from an independent developmental context, these results provide a complementary line of evidence suggesting a link between H2A.V distribution and RNA splicing, and to our knowledge represent the first demonstration of this association in *Drosophila.* Taken together with our transcriptional analysis, this supports a model in which reduced *dom* function influences synaptic communication through coordinated transcriptional and post-transcriptional regulation.

### *Domino* disruption alters synaptic organization in the adult brain

Guided by the transcriptional and splicing changes observed in our analysis, we next examined synaptic organization in adult brains using immunofluorescence staining with the monoclonal antibody nc82, which recognizes the protein Bruchpilot (BRP)(*48*). This protein marks presynaptic active zones and serves as a well-established indicator of synaptic density and organization(*49, 50*). Prominent BRP signal was detected in the antennal lobe, a major olfactory processing center implicated in sleep regulation(*49*) (Fig. 7a, Supplementary Fig. 5e), supporting a potential connection between altered synaptic organization in this region and the sleep phenotypes observed in *dom* heterozygotes. To control for variability across samples, BRP fluorescence intensity was normalized to the background signal measured within the brain foramen, a region expected to lack specific staining. Quantitative analysis revealed a significant increase in BRP fluorescence intensity in both male and female *dom* mutant brains, consistent with altered active zone organization (Fig. 7b, Supplementary Fig. 5f, and Ext. Table 7). In *Drosophila*, sleep deprivation has been shown to elevate BRP expression, linking increased BRP abundance to prolonged wakefulness and heightened synaptic activity(*51*). While our results similarly reveal elevated BRP levels, the underlying sleep phenotypes differ between sexes. This divergence suggests that BRP upregulation in *dom* mutants does not simply reflect increased wakefulness but instead points to broader effects of *dom*-dependent regulation on neuronal structure and function. Collectively, these findings suggest that reduced Domino function influences presynaptic organization and behavior, consistent with a role for chromatin remodeling in shaping neural circuit function.

**Fig. 7:**
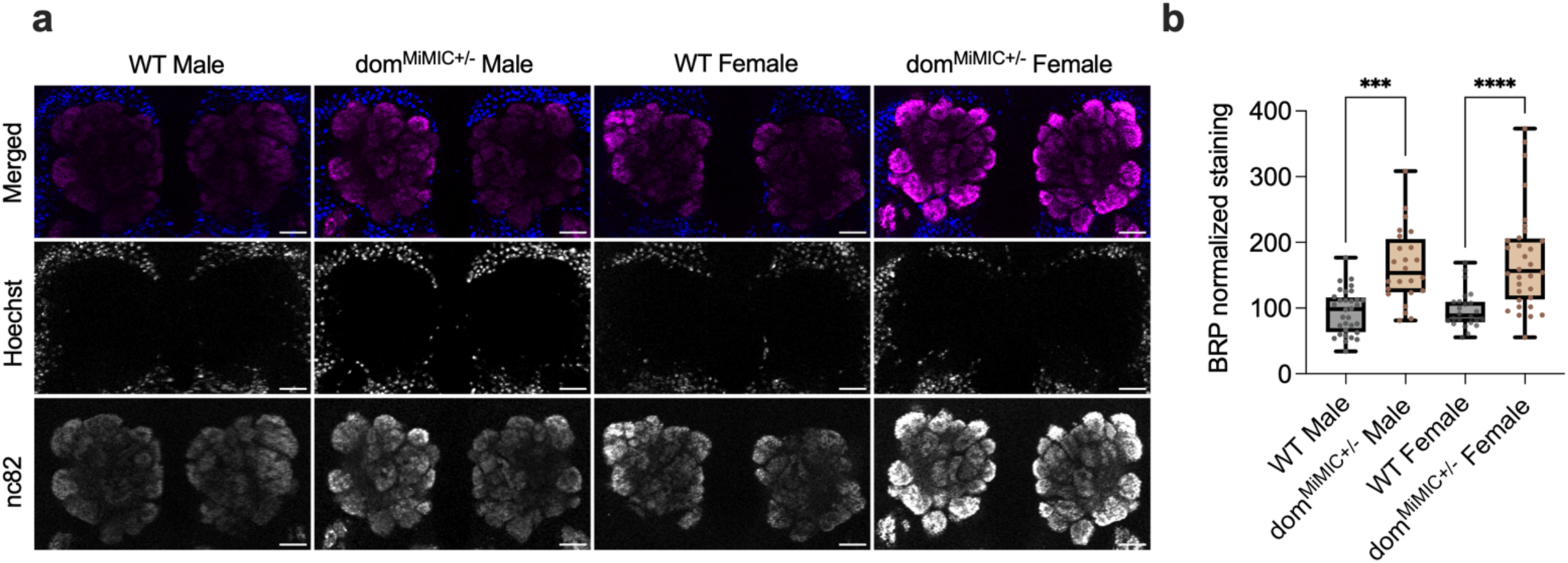
Partial loss of domino alters synaptic organization in dom^MiMIC+/-^ brains. **a**) Immunofluorescence images of antennal lobes stained with Hoechst (blue) and nc82 (magenta), with merged images shown for both sexes. Scale bar 20µm. Brightness and contrast specifications are consistent across images. **b**) Fluorescence intensity was measured as mean ROI antennal lobe - mean ROI foramen, with quantification presented as a percentage of the average wild-type intensity (n=24-32; Two-way ANOVA with Šídák multiple comparisons).

## DISCUSSION

Autism spectrum disorder is thought to arise from the convergence of diverse genetic and molecular perturbations on a limited set of neurodevelopmental pathways. Over the past decade, large-scale human sequencing studies have identified hundreds of ASD-associated genes(*5, 6*). However, for many of these genes, evidence remains primarily statistical and their functional consequences are not yet well defined. To bridge this gap, we turned to *Drosophila melanogaster* as a model system, allowing organism-level evaluation of how individual gene disruptions influence behavior. This strategy enables prioritization of candidate genes based on functional impact and provides an entry point for mechanistic investigation that is not feasible in human cohorts. This study represents one of the most comprehensive efforts to systematically link high-confidence ASD-associated genes to behavioral phenotypes in a living organism. To enable screening at this scale, we refined the analysis of an established activity assay(*14, 17*) for comprehensive multi-parameter profiling and developed FlyFinder, an automated image analysis pipeline that quantifies social spacing from group assays(*21, 22*). These methodical advances allowed us to efficiently interrogate dozens of gene perturbations while maintaining the sensitivity needed to detect subtle behavioral differences.

While no single assay can capture the full complexity of ASD-relevant phenotypes, we selected complementary measures that quantify conserved dimensions of behavior. The activity assay measures alterations in activity patterns and sleep architecture rather than directly modeling specific human behaviors, but it provides reproducible and quantifiable readouts of behavioral state. Because this assay yields multiple interrelated parameters describing movement and rest, we used PCA to summarize these measures and identify shared patterns across genotypes. However, PCA necessarily simplifies complex behavioral variance and may obscure gene-specific effects by compressing distinct behavioral changes into shared summary dimensions. Therefore, interpretation of behavioral outcomes can integrate both component-level trends and individual assay results to ensure robustness.

Social spacing provides a quantitative measure of the preferred inter-individual distance maintained in a group and reflects how flies perceive and respond to one another. Importantly, the direction of change does not determine ASD relevance; both increased clustering and increased spacing represent deviations from typical group spacing behavior. Consistent with the genetic heterogeneity of ASD, different risk-gene orthologs shifted social spacing in opposite directions (e.g., *dom* mutants decreased whereas *kn* mutants increased spacing), suggesting that distinct perturbations influence social behavior through different mechanisms.

Integrating social spacing with sleep and activity measures provides a stronger basis for prioritizing genes and designing follow-up experiments. While the observed behaviors do not model ASD per se, they capture dimensions of neural regulation that are frequently disrupted across neurodevelopmental disorders. The high proportion of behavioral hits indicates that many ASD risk genes exert conserved, dosage-sensitive effects on neural circuits, and that partial loss of gene function alone is sufficient to induce behavioral changes. Among the genes identified through this approach, the chromatin remodeler *domino* (*dom*), orthologous to human *Snf2-related CREBBP activator protein (SRCAP)*, emerged as a regulator of each behavioral domain. *dom* encodes a chromatin remodeler mediating the incorporation of histone variant H2A.V into nucleosomes, which shapes transcriptional accessibility(*28, 52, 53*). In mammals and yeast, H2A.Z (orthologous to *Drosophila* H2A.V) has been shown to be essential for splicing integrity and for regulating genes involved in neuronal structure and synaptic function(*44, 45, 54*). Here, we extend this paradigm to *Drosophila* by showing that *dom* heterozygosity disrupts transcriptional programs in adult brains. While changes in male and female brains differ, they converge on pathways associated with synaptic signaling, consistent with shared behavioral phenotypes. Further analyses revealed widespread changes in RNA splicing in the brains of both dom^MiMIC+/–^ and dom^Trojan+/–^ flies. These splicing alterations affected all major event types and were enriched for genes involved in synaptic signaling, neuronal maturation, and developmental processes, underscoring the broad regulatory impact of partial *dom* loss. Reanalysis of a published embryo dataset(*47*) further demonstrated that *dom* knockdown alters H2A.V distribution, with a global reduction at transcription start and end sites and a redistribution of H2A.V over the bodies of differentially spliced genes. To our knowledge, this represents the first evidence in *Drosophila* linking H2A.V localization to splicing outcomes, extending findings from mammalian and yeast systems. Together, the widespread transcriptional and splicing alterations in brains of *dom* heterozygote animals link *dom* to transcriptional and post-transcriptional regulation that supports proper synaptic function and neural communication. This model aligns with a growing body of evidence implicating chromatin regulators in ASD pathogenesis, including chromodomain helicase DNA-binding (CHD)(*55, 56*) family of chromatin remodeling enzymes and ARID1B(*57*), which similarly influence transcriptional programs governing neuronal connectivity.

These molecular and post-transcriptional changes align closely with the structural phenotypes we observed. Both dom^MiMIC+/–^ and dom^Trojan+/–^ flies exhibited increased Bruchpilot (BRP) signal in the antennal lobe, indicating elevated active-zone density. Because the antennal lobe integrates sensory and neuromodulatory inputs relevant to sleep and arousal(*49*), altered synaptic organization in this region provides a compelling link between the molecular consequences of Domino loss and the behavioral phenotypes identified in our screen.

Our analyses also revealed pronounced sex differences in behavioral and transcriptional responses, while the structural changes were evident in both sexes. Sex-biased effects are increasingly recognized in neurodevelopmental disorders(*58*) and may reflect differences in hormonal signaling or neural circuit modulation(*41–43*). Consistent with this, our transcriptomic data identified enrichment of hormone receptor and signaling pathways across sexes, supporting a potential interaction between chromatin regulation and hormonal state in shaping behavioral outcomes.

From an evolutionary and translational perspective, the conservation of SRCAP function underscores the relevance of *Drosophila* for modeling chromatin-based mechanisms of neurodevelopmental disorders. Our cell-specific knockdown experiments show that *dom* is required across neural and non-neural cell types. Consistent with this essential role, our immunofluorescence analysis revealed altered synaptic zone organization in *dom* heterozygous mutant brains, providing structural evidence that disruptions in chromatin remodeling impact synaptic architecture. The behavioral and molecular parallels between *dom* mutants and *SRCAP*-associated disorders(*29, 30, 59*) suggest that partial reduction of Domino/SRCAP activity disrupts a conserved transcriptional-splicing program essential for synaptic homeostasis.

Together, these findings support a model in which chromatin remodeling by Domino/SRCAP coordinates transcriptional and post-transcriptional regulation to maintain synaptic structure and function, thereby influencing ASD-relevant behaviors. More broadly, this work demonstrates how combining behavioral and transcriptomic analyses can uncover molecular mechanisms underlying complex neurodevelopmental traits. Future studies will be needed to determine how Domino-dependent chromatin regulation is interpreted across specific neuronal populations and circuits, and whether similar programs are engaged by other ASD-associated chromatin regulators.

## MATERIALS AND METHODS

### Fly Husbandry

Fly stocks were maintained on standard organic diet purchased from LabExpress (inactivated yeast 16 g/L, organic soy corn flour 9 g/L, organic yellow cornmeal 67 g/L, agar 6 g/L, organic light corn syrup 71 mL/L, propionic acid 4.4 mL/L) at 25°C with 60% humidity on a 12 h light-dark cycle. Fresh vials and bottles were regularly seeded with adult flies (< 7 days old) and maintained at consistent population densities. To prevent interaction between parental and progeny generations, adult flies from the previous cohort were removed prior to eclosion of the next generation.

Stocks obtained from the Bloomington Drosophila Stock Center (NIH P40OD018537) were used in this study: MiMIC^+/-^ lines (Ext. Table 1), dom^MiMIC+/-^ (RRID:BDSC_44733), dom^Trojan+/-^ (RRID:BDSC_76192), TRiP line control (RRID:BDSC_36303), *dom* RNAi (RRID:BDSC_34827), tubulin-Gal4 (RRID:BDSC_5138), elav-Gal4 (RRID:BDSC_458), repo-Gal4 (RRID:BDSC_7415), MB-Gal4 (RRID:BDSC_4440). Our wild-type strain (Harwich 15) was a gift from T. Eickbush. We used FlyBase(*60*) (FB2021_06) to obtain information on the MiMIC insertion orientations for strains used within the screen, while subsequent versions through FB2025_05 were consulted for phenotypes/function/stocks/gene expression (etc).

### Screen Experimental Design

Virgin wild-type females were collected and maintained in cages on a standard sugar-yeast diet (glucose 15 g/L, yeast 100 g/L, agar 15 g/L, propionic acid 6 mL/L, nipagin 15 mL/L) for 10 days. Females were then mated with either MiMIC mutant males or wild-type controls, and males were removed after 6 h of mating. All MiMIC lines were derived from the MiMIC gene disruption collection maintained in a y^1^ w* background(*15*), resulting in white-eyed flies. Because mutations in *white* have been reported to potentially influence visual function and affect behavioral assays, particularly those involving social interaction(*21*), mutant males were crossed to red-eyed Harwich wild-type females prior to testing.

The following morning, 100 µL of embryos suspended in PBS were collected and transferred to sugar-yeast vials (8-10 vials per condition) and allowed to develop under standard conditions. Behavioral traits in *Drosophila* exhibit substantial inter-individual variability and are sensitive to environmental and genetic background effects. To minimize these sources of variance, all lines were reared under standardized density, temperature, and light–dark conditions. Newly eclosed offspring (1-2 days post-eclosion) were collected and maintained on fresh sugar-yeast food for 3-4 days prior to phenotypic testing. Behavioral assays were conducted on red-eyed progeny, ensuring consistent background contribution across lines. Behavioral measurements were performed with sufficient sample sizes, and each phenotype was assessed across 2-3 independent biological replicates to ensure reproducibility. Wild-type Harwich flies were assayed in parallel within each experimental batch as negative controls.

For X-linked insertions, male progeny inherited the wild-type maternal X chromosome and therefore did not carry the mutation and were excluded from analysis. Although no designated positive-control mutant were included across all assays, several phenotypes observed in the screen were consistent with previously reported behavioral effects for select ASD-associated genes (*brm*(*18*)*, bchs*(*19*)*, Nlg3*(*22*)), providing internal validation of assay sensitivity. Phenotypes that were inconsistent across biological replicates or that did not segregate with the targeted insertion were interpreted cautiously as potential background or environmental effects.

### Activity Assay

Flies were individually placed into Trikinetics DAM2 monitors with a standard sugar-yeast food source and allowed to acclimate for approximately 10 h before the start of the recording period. Locomotor activity was then monitored in 1 min intervals for the next 48 h. Data were initially processed using the *Drosophila* Sleep and Circadian Analysis MATLAB Program (SCAMP)(*17*). The 51 output measures generated by SCAMP were subjected to PCA using the prcomp function in R v4.4.1, with analyses stratified by sex. Prior to PCA, measures were log2 transformed and where necessary and a constant was added to shift values above zero before transformation. Linear mixed-effects (LME) models were fit for each PC using the nlme package(*61*). Variances modelled as varying by genotype using the varIdent weights and random effects were included for each replicate (3-4 flies per breeding vial) nested within experimental date.

For the top 6 PCs, we estimated 95% confidence intervals for two comparisons: (i) each mutant genotype versus wild-type, and (ii) each genotype versus the pooled mean of all other mutant lines (e.g., *dom* compared with the mean of all non-*dom* and non-wild-type genotypes). Two SGPVs(*62*) were computed for each comparison, each using a ±10% null interval. The first SGPV tested whether each genotype differed from wild-type; the second tested whether a genotype differed from all other mutant lines. A genotype was considered significant only when both criteria were met (SGPV = 0).

Results were visualized using custom rose plots (generated with ggplot2(*63*) and adapted from circular barplots(*64*)), forest plots (estimates with 95% confidence intervals colored by significance), and beeswarm plots of mutant minus wild-type estimates(*65*).

To ensure comparability between the *domino* validation lines and the full-screen dataset, PCA for both dom^MiMIC+/-^ and dom^Trojan+/-^ lines used the same rotation matrix derived from the full screen. The first 6 PCs of each line were analyzed using the same regression models as the full screen. Because these analyses constituted validation assays, statistical significance was assessed using nominal p-values (p < 0.01 was considered significant).

### Social Spacing Assay

The social spacing assay was adapted for high-throughput use from previously described methods(*21*). Fly vials containing mixed-sex populations were acclimated to the behavioral room conditions for at least 2 h prior to testing. All experiments were conducted between zeitgeber time (ZT) 5 and ZT 9 to control for circadian effects. Flies were briefly anesthetized with CO_2_ and separated by sex into groups of 20 before being placed in a vertical triangular arena (height = 13 cm, base = 13 cm). The four triangle panels were arranged in a square grid, allowing simultaneous testing of multiple experimental conditions. Once all flies had recovered from anesthesia, they were gently tapped to the bottom of the arena to ensure a uniform starting position. After 20 min of exploration, when flies had settled into their preferred social spacing, an image was captured and subsequently processed using FlyFinder (https://github.com/LempradlLab/drosophila_screen_dom).

FlyFinder, a custom Python-based image segmentation and analysis pipeline, was developed to automate the identification and spatial analysis of individual *Drosophila* within the multi-arena images. The program uses a random forest classifier, trained on 100 manually labeled flies, to distinguish flies based on size and pixel density. Each image was segmented to isolate a single arena, and individual flies were identified within each triangular arena. Manual verification and correction of segmentation errors were performed in Fiji(*66*), followed by a secondary segmentation pass to ensure accuracy. The program extracts x-y coordinates for all identified flies and calculates multiple spatial and behavioral parameters from these coordinates (number of flies within 4 body lengths, 3 nearest neighbors, nearest neighbor, and global median). Developed in Python and R, FlyFinder includes Jupyter Notebooks and Google Colab workflows for streamlined, reproducible analysis.

FlyFinder outputs were analyzed using either log₂-transformed outcomes (3 nearest neighbors, nearest neighbor, and global median) or negative binomial regression for count outcomes (number of flies within 4 body lengths). Log-transformed outcomes were modeled using LME models (nlme)(*61*), specifying unequal variances across genotypes and random effects for each replicate of 20 flies nested within experimental date. Count outcomes were modeled using glmmTMB, the *nbinom2* family, and the same random-effects structure(*67, 68*).

Outcomes that could take negative values were shifted to be strictly positive by adding 1 plus the absolute value of the minimum observed value for that outcome. Analyses were stratified by sex. For each outcome, we calculated the same two SGPVs used in the activity analysis, (i) comparing the genotype with wild-type and (ii) comparing genotype with all other mutant lines. Genes were considered hits if they exhibited SGPVs indicating meaningful differences from both wild-type and all other mutant genotypes (SGPV = 0). Results were visualized as custom forest plots and radar plots generated with ggplot2 and ggradar(*63, 69*). For the screen radar plots, a t-statistic (estimate divided by standard error) was used from the mutant vs wild-type contrast for each behavioral measure. T-statistics were shifted to a non-negative scale so that a value of zero, indicating no difference from wild-type, corresponds to the inner reference polygon. A single shared axis scale was applied across all genotypes and both sexes to allow direct comparison of effect sizes, with the outer ring corresponding to the largest t-statistic observed across the dataset. The dom^MiMIC+/-^ and dom^Trojan+/-^ validation lines were each analyzed using the same regression models, with radar plots generated using the same approach: male and female plots share a common axis scale to allow direct comparison, with statistical significance based on the nominal p-value being less than 0.05.

### Rapid Iterative Negative Geotaxis (RING) Assay

At least 24 h prior to behavioral testing, flies were sorted by sex and transferred to fresh vials containing a standard sugar-yeast diet, with 10 flies housed per vial. Vials were maintained at 25°C and 60% relative humidity. Immediately before testing, flies were transferred, without anesthesia, into empty plastic vials and placed within the TriKinetics LAM25 monitoring system attached to a standard multi-tube vortexer (Talboys). Over a 10 min testing period, flies were subjected to 3 s of vigorous shaking (4^th^ intensity setting) at 30 s intervals. Activity readings were collected every 5 s to quantify the number of flies crossing a 2 cm threshold. The average number of climbs per fly was calculated for each 5 s interval, excluding the initial measurement taken within the first 5 s following the onset of shaking. Statistical analyses were conducted using Two-way ANOVA with Šídák multiple comparisons for males and females.

### RNAi

Crosses were established using 5 virgin Gal-4 driver females and 2 males from either the TRiP control or the *dom* RNAi lines. Flies were allowed to mate for 2 days before the vials were left undisturbed for development through eclosion. Pupal numbers were recorded from counts along the vial walls, and adults were scored post-eclosion for the appropriate phenotype (balance versus non-balancer). Statistical comparisons were performed using the nonparametric Mann-Whitney test.

### RNA Extraction

Prior to RNA extraction, brain samples (5 brains per replicate) were dissected in 0.1% PBST (Tween 20, Bio-Rad, 1706531) and immediately frozen with liquid nitrogen in screw cap tubes containing ∼0.2 g Lysing Matrix D beads (MP Biomedicals, 6540434) and stored at -80°C. 500 uL TRIzol Reagent (Life Technologies, 15596026) was added to each tube, then homogenized using the FastPrep-24 system on high for 30 s and visually observed for tissue disruption. 100 μL chloroform (Sigma-Aldrich, 319988) was added and samples were centrifuged at 12,000 x g for 15 min at 4°C. The upper aqueous phase was transferred to new Eppendorf tubes followed by RNA precipitation using 250 μL ice-cold isopropanol (Sigma-Aldrich, 190764) and 1 μL GlycoBlue (ThermoFisher, AM9516) for pellet visualization. Samples were centrifuged at 12,000 x g for 15 min at 4°C. Supernatant was removed and discarded. The samples were washed twice with 75% ethanol (Fisher Scientific, 111000200), using a 5-min 7,500 x g centrifugation each round for pellet collection. The pellet was air dried for 10 min and resuspended in 10 μL RNase-free water (Life Technologies, AM9938). Samples were incubated at 60°C for 5 min, then stored at -80°C.

### Quantitative PCR (qPCR)

Adult fly heads were collected, flash-frozen in liquid nitrogen, and stored at - 80°C until processing. Samples were homogenized in TRIzol Reagent (Life Technologies, 15596026) and total RNA was isolated as described above, with RNA purity and concentration assessed via NanoDrop spectrophotometer. Residual genomic DNA was removed using the TURBO DNA-free kit (Thermo Fisher, AM1907) following the manufacturer’s instructions. Complementary DNA (cDNA) was synthesized from 1 μg of total RNA using an oligo(dT)_12-18_ primer (Thermo Fisher, 18418012) and the M-MLV Reverse Transcriptase kit (Promega, M1701). Quantitative PCR (qPCR) reactions were performed using SsoAdvanced Universal SYBR Green Supermix (BioRad, 1725271). Thermal cycling conditions consisted of an initial denaturation at 98°C for 30 s, followed by 45 cycles of 98°C for 10 s and 58°C for 30 s. A melt-curve analysis was included to confirm amplification specificity. Each sample was analyzed with 6 technical replicates, and no-template controls were included to confirm the absence of contamination. Relative gene expression was calculated using the ΔΔCt method, with normalization to the reference gene *Tubulin*. Statistical analyses were performed using a Kruskal-Wallis test for nonparametric data, with multiple comparison tests conducted separately for the dom^MiMIC+/-^ and dom^Trojan+/*-*^ lines.

Primers were designed using NCBI Primer-BLAST to span exon-exon junctions and avoid amplification of genomic DNA. *dom* forward primer: CAGCTAACGCAACAAGGTGG, *dom* reverse primer: TCCGCTTGAGGCATTGTTCT, *tubulin* forward primer: TGTCGCGTGTGAAACACTTC, *tubulin* reverse primer: AGCAGGCGTTTCCAATCTG.

### Construction and Sequencing of Directional total RNA-seq Libraries

Libraries were prepared by the Van Andel Genomics Core from 500 ng of total RNA using the KAPA RNA HyperPrep Kit (Kapa Biosystems, Wilmington, MA USA). Ribosomal RNA material was reduced using the QIAseq FastSelect –rRNA Fly Kit (Qiagen, Germantown, MD, USA). RNA was sheared to 300-400 bp. Prior to PCR amplification, cDNA fragments were ligated to IDT for Illumina TruSeq UD Indexed adapters (Illumina Inc, San Diego CA, USA). Quality and quantity of the finished libraries were assessed using a combination of Agilent DNA High Sensitivity chip (Agilent Technologies, Inc.), QuantiFluor® dsDNA System (Promega Corp., Madison, WI, USA), and Kapa Illumina Library Quantification qPCR assays (Kapa Biosystems). Individually indexed libraries were pooled and 100 bp, paired-end sequencing was performed on an Illumina NovaSeq6000 sequencer to an average depth of 15M raw paired-reads per transcriptome. Base calling was done by Illumina RTA3 and the output of NCS was demultiplexed and converted to FastQ format with Illumina Bcl2fastq v1.9.0.

### ChIP-Seq Analysis

Reads were processed with the VARI-BBC pipeline (https://github.com/vari-bbc/Peaks_workflow). Briefly, reads were trimmed using TrimGalore v0.6.10 (https://github.com/FelixKrueger/TrimGalore) with default settings. Trimmed reads were aligned to the *D. melanogaster* reference genome dm6_BDGP6.28.100 using bwa mem v0.7.17(*70*). Alignments were filtered with samtools view v1.17(*71*) with the parameters ‘-q 30 -F 2828’ and ‘-f 2’ for paired-end reads. The csaw package v.1.38.0 was used to find peaks(*72*). The windowCounts function was used to find peaks using the parameters, ‘width = 10000, minq = 20, dedup = TRUE, restrict = c("2L", "2R", "3L", "3R", "4", "X", "Y")’ and using ENCODE’s dm6_BDGP6.28.100 blacklist. Efficiency biases were removed by creating a second set of high-abundance peaks with the windowCounts function with ‘width = 150’, and the same parameters. The filterWindowsGlobal function was used to filter windows, which were then used to normalize the peaks using the normFactors function and subsequently scaled with the scaledAverage function to find the final peaks. Differential binding was done with edgeR’s (v.4.0.16) glmQLFit function with the ‘robust = TRUE’ parameter(*73*).

### Gene Profile Plots

Gene profile plots were created by first combining sample bigwigs generated by the aforementioned VARI-BBC workflow into group wig files using wiggletools mean(*74*), and converted back to bigwigs with wigToBigWig(*75*). BED files containing genetic loci of significantly differentially spliced (fdr < 0.05) genes and an equal number of near-significantly spliced genes (fdr > 0.05) were acquired from the dm6_BDGP6.28.100 annotation GTF. BigWig and BED files were then used with deeptools computeMatrix(*76*) tool along with the dm6_BDGP6.28.100 blacklist, and parameters: ‘-startLabel "TSS", -endLabel "TES", -b 200, -a 200, -p 8, --smartLabels’ to build the plot matrix. The matrix can be passed to deeptool’s plotProfile function or used to build custom plots.

### Differential Expression, Overrepresentation, Gene Set Enrichment, and Splicing Analyses

Low quality read ends were trimmed and adaptor sequence were removed using Trimgalore (https://github.com/FelixKrueger/TrimGalore) v0.6.0 (dom^MiMIC+/-^ analysis) and Trimgalore v0.6.1 (dom^Trojan+/-^ analysis) with --quality parameters set to 20. Trimmed reads were aligned to the *Drosophila melanogaster* reference genome (dm6_BDGP6.28.100) using STAR v2.7.8a (dom^MiMIC+/-^ analysis) and STAR v2.7.11b (dom^Trojan+/-^ analysis) with default parameters. Aligned reads were quantified with featureCounts (-M --fraction) to include multimapping reads. Fractional assignments were rounded to the nearest integer prior to downstream analyses. The resulting multimapping .txt files were imported into R v4.4.1 to generate raw counts tables. Differential expression analysis was conducted using DESeq2 v.1.46.0, and differentially expressed (DE) genes are provided in Ext. Table 4. ORA of shared significant genes between males and females (padj ≤ 0.01 and |log2fold change| ≥ 0.5) was performed using FlyEnrichr(*39, 40*) (padj < 0.05). GSEA(*36, 37*) was performed using normalized count data and visualized in Cytoscape(*38*) (Q-value 0.25, Edge Cutoff 0.25 Jaccard Index). Differential splicing analysis was conducted using rMATS(*46*) with STAR alignments. Significant splicing events were filtered from the .JCEC output files based on false discovery rate (10 count threshold, FDR ≤ 0.05, and InclusionLevelDifference ≤ 0.1). ORA of significant splicing events was also performed using FlyEnrichr(*39, 40*) (padj < 0.01).

### Immunostaining

Adult flies were fixed with 4% Formaldehyde (Sigma-Aldrich, F8775) in 0.3% PBSX (Triton-X, Fisher Scientific, BP151) overnight at 4°C. Brains were dissected in 0.1% PBST (Tween 20, Bio-Rad, 1706531), permeabilized in 0.3% SDOC (Alfa Aesar, B20759), washed 3x in PBSTX (0.1% Tween 0.3% Triton-X), blocked in 5% BSA (Fisher Scientific, BP9700) and incubated in primary antibody (anti-nc82 1:500, DSHB, AB_2314866 concentrate) in 3% BSA overnight at 4°C. Samples were washed in 300 mM NaCl (Millipore, 7647-14-5) and 3x with PBSTX, then incubated in the secondary antibody (anti-mouse 647 1:1000, Life Technologies, A21236) in 3% BSA for 1.5 h at RT. Finally, samples were washed with 300 mM NaCl and 3x with PBSTX, incubated with Hoechst 33342 (Thermo Scientific, 62249) for 10 min at RT and washed again 3x with PBSTX. Samples were mounted on bridge slides and stored at 4°C.

The monoclonal antibody nc82, developed by Buchner, E., was obtained from the Developmental Studies Hybridoma Bank, created by the NICHD of the NIH and maintained at The University of Iowa, Department of Biology, Iowa City, IA 52242.

Slides were imaged on the Zeiss Axio Observer 7 with Apotome at 20X magnification. Z-stacks were acquired at 4 μm intervals to capture the full depth of the brain. Images were analyzed using Fiji(*66*), ensuring consistent imaging parameters and processing settings across all conditions. For each sample, the slice with the highest anatomical resolution of the antennal lobe was identified, and three slices above and three slices below this plane were selected to generate a seven-slice *z*-projection using the average-intensity. Quantification was restricted to the antennal lobe region. Fluorescence intensity was measured as mean ROI antennal lobe - mean background ROI central foramen, presented as a percentage of the average wild-type intensity (Two-way ANOVA with Šídák multiple comparisons). Samples were excluded from analysis if tissue obstructed the foramen or if the boundaries of the antennal lobe could not be clearly delineated (Ext. Table 7; red text).

## Supporting information

Ext. Table 1

Ext. Table 2

Ext. Table 3

Ext. Table 4

Ext. Table 5

Ext. Table 6

Ext. Table 7

## Data Availability

RNA-seq data generated in this study have been deposited under BioProject accession PRJNA1401847 and SRA accession SUB15928845. All custom scripts used for data processing, analysis, and Fig. generation are available at https://github.com/LempradlLab/drosophila_screen_dom).

## Author Contributions

Conceived and designed the experiments: ES AL. Performed the experiments: ES JR DM. Analyzed the data: ES ZM BM RF DC SV AL. Wrote the paper: ES AL.

## Acknowledgements

This work was supported by a MeNu Pilot Award from Van Andel Institute – Metabolism and Nutrition (MeNu) Program (RRID:SCR_027494). We also thank the Van Andel Institute Bioinformatics and Biostatistics Core (RRID:SCR_024762), Genomics Core (RRID:SCR_022913) and the Optical Imaging Core (RRID:SCR_021968) for their assistance with this work. We are grateful to all members of the Lempradl Lab for their valuable discussion and feedback throughout this project, as well as their thoughtful comments on the manuscript. Generative AI tools (ChatGPT 5.2) were used to assist with language editing and clarity of the manuscript. All scientific content, analyses, and interpretations were developed and verified by the authors.

---

**Supplementary Fig. 1:**
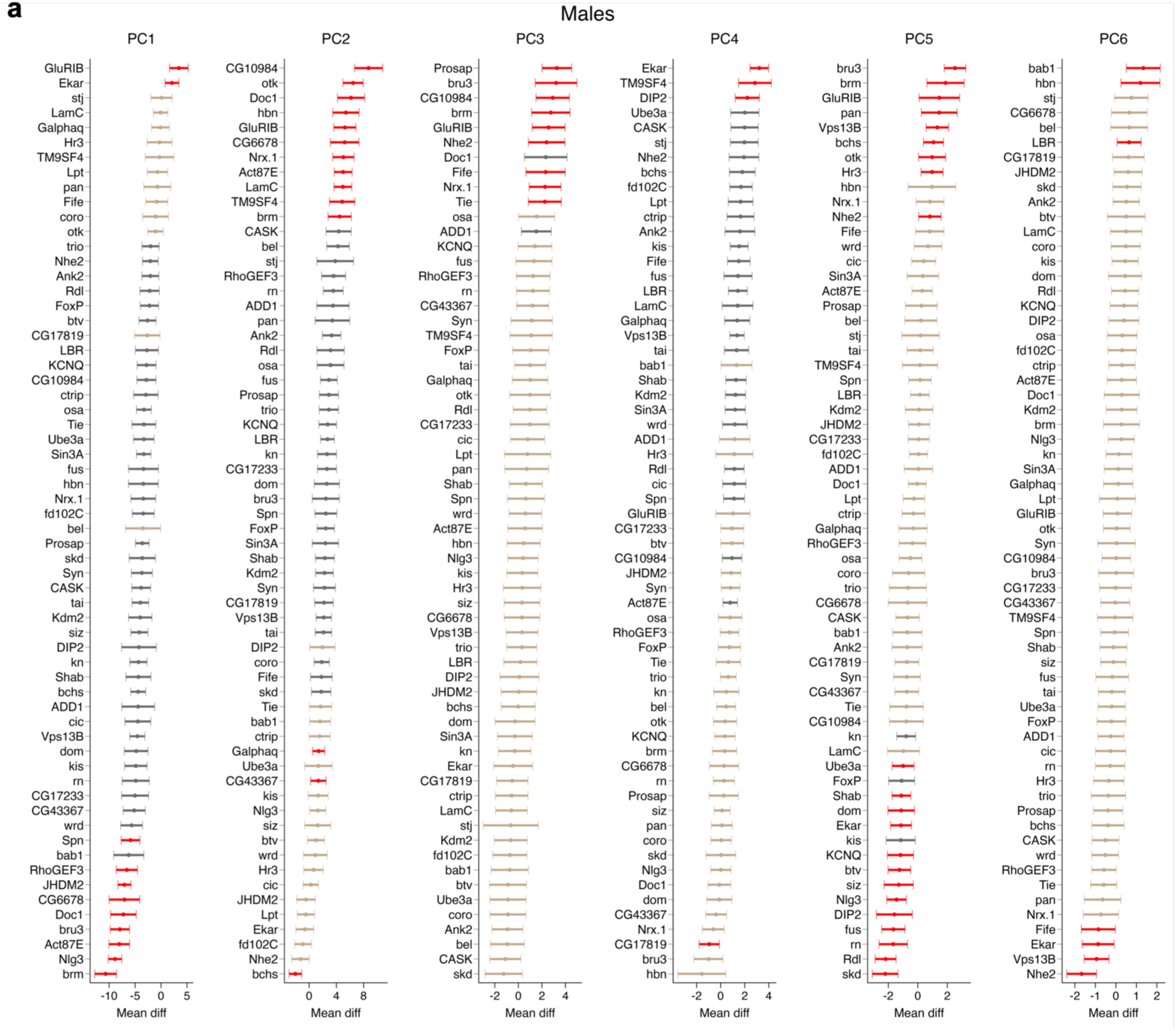
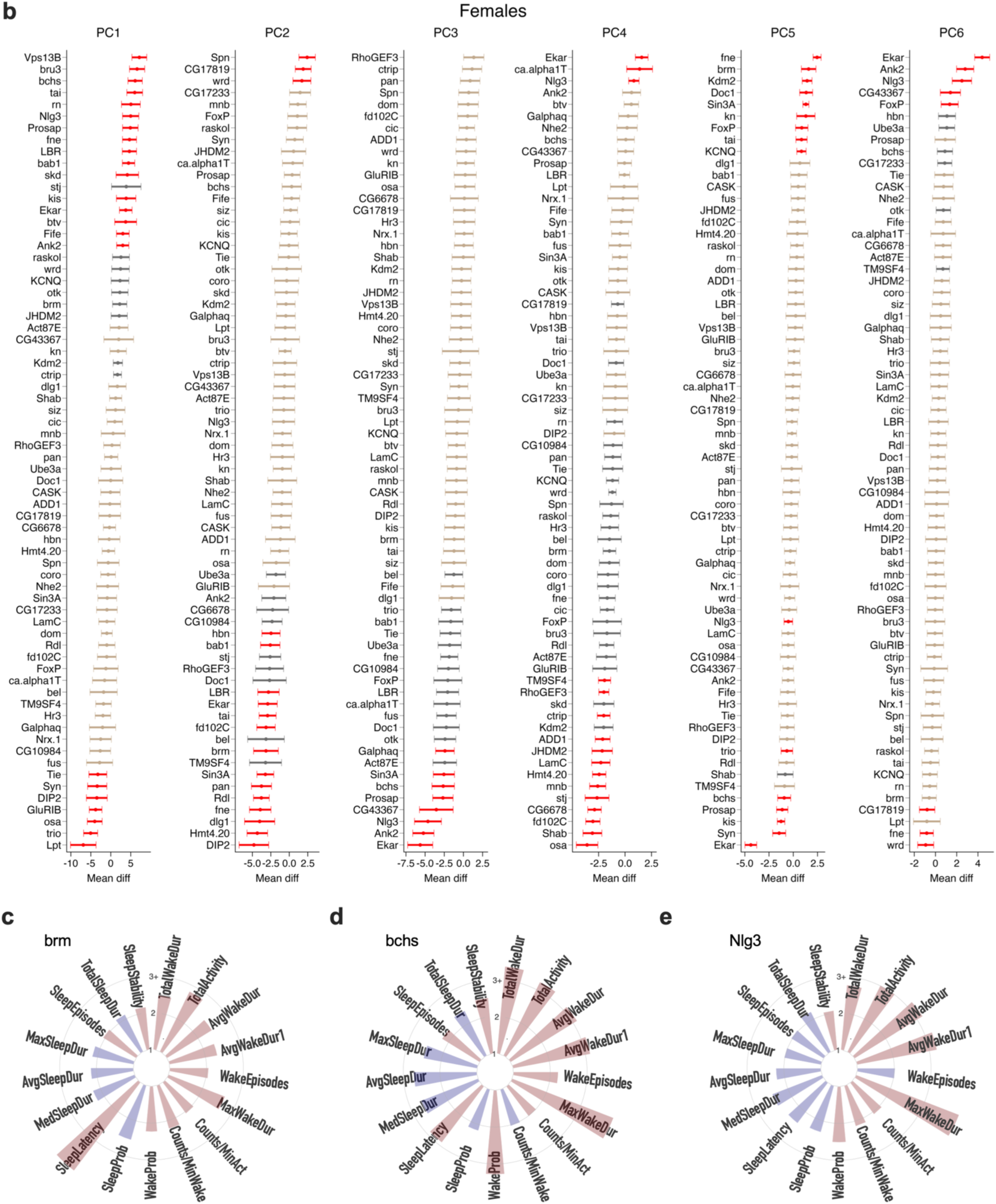
Genetic effects on sleep and activity phenotypes. **a,b**) Forest plots for all 68 viable screened genes in males (**a**) and females (**b**) (n=2-3 replicates per line with 4 flies per replicate). For the top 6 PCs, 95% confidence intervals were computed for two comparisons: (i) each mutant genotype versus wild-type and (ii) each genotype versus the pooled mean of all other mutant lines. SGPVs were calculated for each comparison. Genes are ranked within each PC by estimated effect size, with confidence intervals shown. Genes meeting both hit criteria (SGPV = 0) are shown in red, those meeting only the wild-type comparison are shown in gray, and inconclusive genes are shown in beige. **c-e**) Rose plots of top genes significantly different by both criteria and shared between sexes, shown for females. Plots are organized clockwise from “more awake” to“more asleep” and summarized across the full 24 h monitoring period. Red bars indicate increased behavior in MiMIC^+/-^ flies, while blue bars indicate decreased behavior.

**Supplementary Fig. 2:**
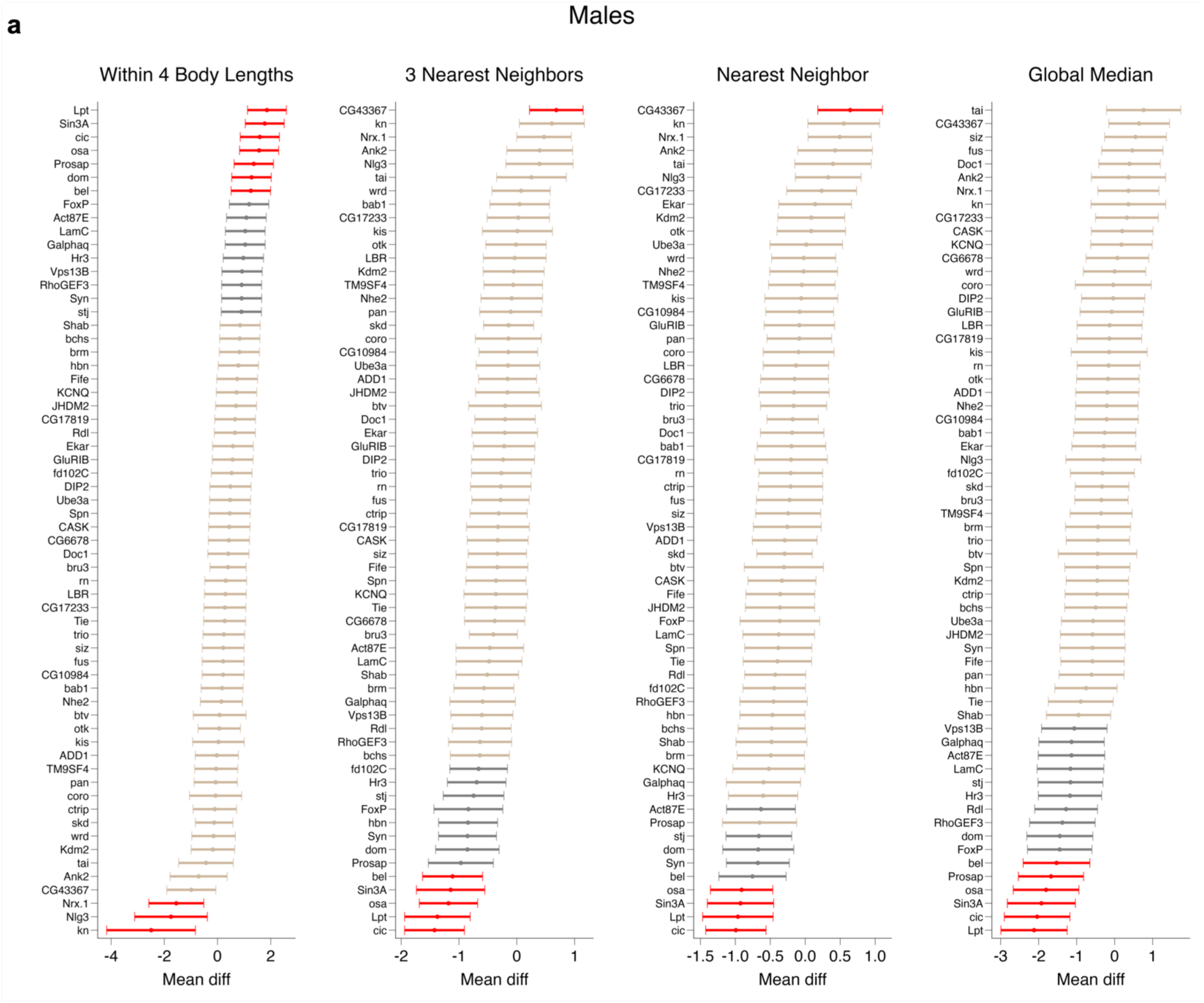
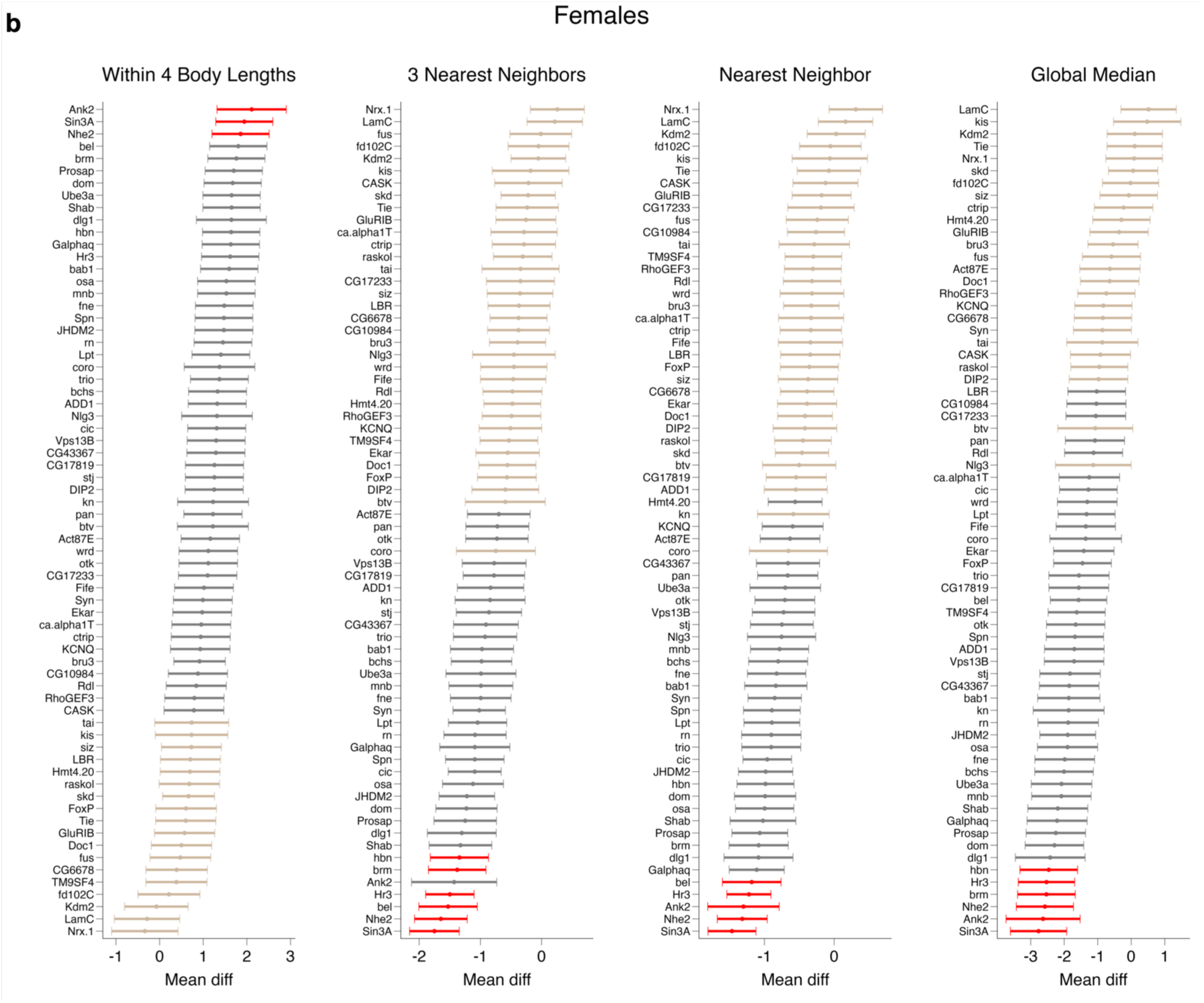
Genetic effects on social behavior phenotypes. **a,b**) Forest plots for all 68 viable screened genes in males (**a**) and females (**b**) (n=2-3 replicates per line with 4 flies per replicate). SGPVs were calculated for the comparisons (i) genotype vs wild-type and (ii) genotype vs all other mutant lines. Genes are ranked within each social spacing parameter by estimated effect size, with confidence intervals shown. Genes meeting both hit criteria (SGPV = 0) are shown in red, those meeting only the wild-type comparison are shown in gray, and inconclusive genes are shown in beige.

**Supplementary Fig. 3:**
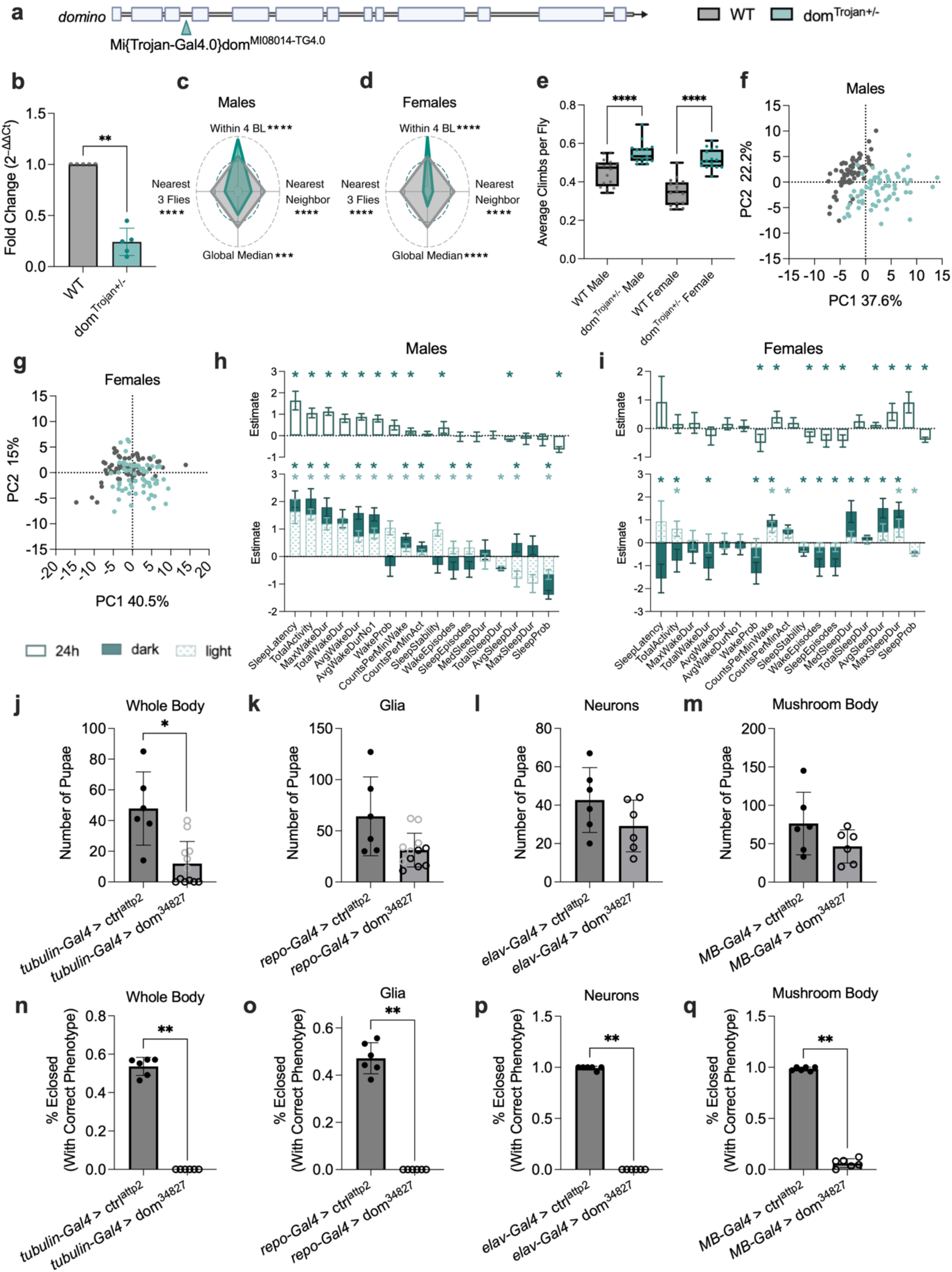
Behavioral characterization of dom^Trojan+/-^ flies. **a**) Schematic of the domino^Trojan+/-^ insertion. **b**) qPCR analysis of dom expression in wild-type and dom^Trojan+/-^ flies. (n=5; 10 heads per replicate; Kruskal-Wallis test). **c,d**) Radar plots of social spacing assay results for males (**c**) and females (**d**), showing t-statistics from the *dom* vs wild-type contrasts (n=16; 20 flies per replicate). The inner polygon represents the wild-type reference (t = 0), with axes extending outward indicating higher values relative to wild-type and axes plotting inward indicating lower values. Male and female plots share a common axis scale to allow for direct comparison. **e**) Climbing assay performance in males and females (n=16; Two-way ANOVA with Šídák multiple comparisons). **f,g**) PCA of activity assay results in males (**f**) and females (**g**), with wild-type (gray) and dom^Trojan+/-^ (teal) flies (n=64). **h,i**) Individual sleep and activity parameters for males (**h**) and females (**i**). For each measure, the 24 h estimate is shown in the upper graph (open bars), with light- and dark-period estimates shown in the lower graph (patterned and filled bars, respectively). Estimates are plotted such that positive values indicate an increase and negative values indicate a decrease in dom^MiMIC+/-^ flies relative to controls. Asterisks indicate significance (p ≤ 0.01), with light and dark blue asterisks denoting significance during the light and dark periods, respectively. **j-m**) Total pupal counts from *dom* RNAi crosses targeting (**j**) tubulin-Gal4 (whole-body), (**k**) repo-Gal4 (glia), (**l**) elav-Gal4 (neurons) and (**m**) MB-Gal4 (mushroom body). Light gray open circles represent pupae that were wild-type due to the presence of a balancer chromosome. **n-q**) Percentage of total pupae that successfully eclosed to adults for the same RNAi crosses shown in **j-m** (Mann-Whitney test).

**Supplementary Fig. 4:**
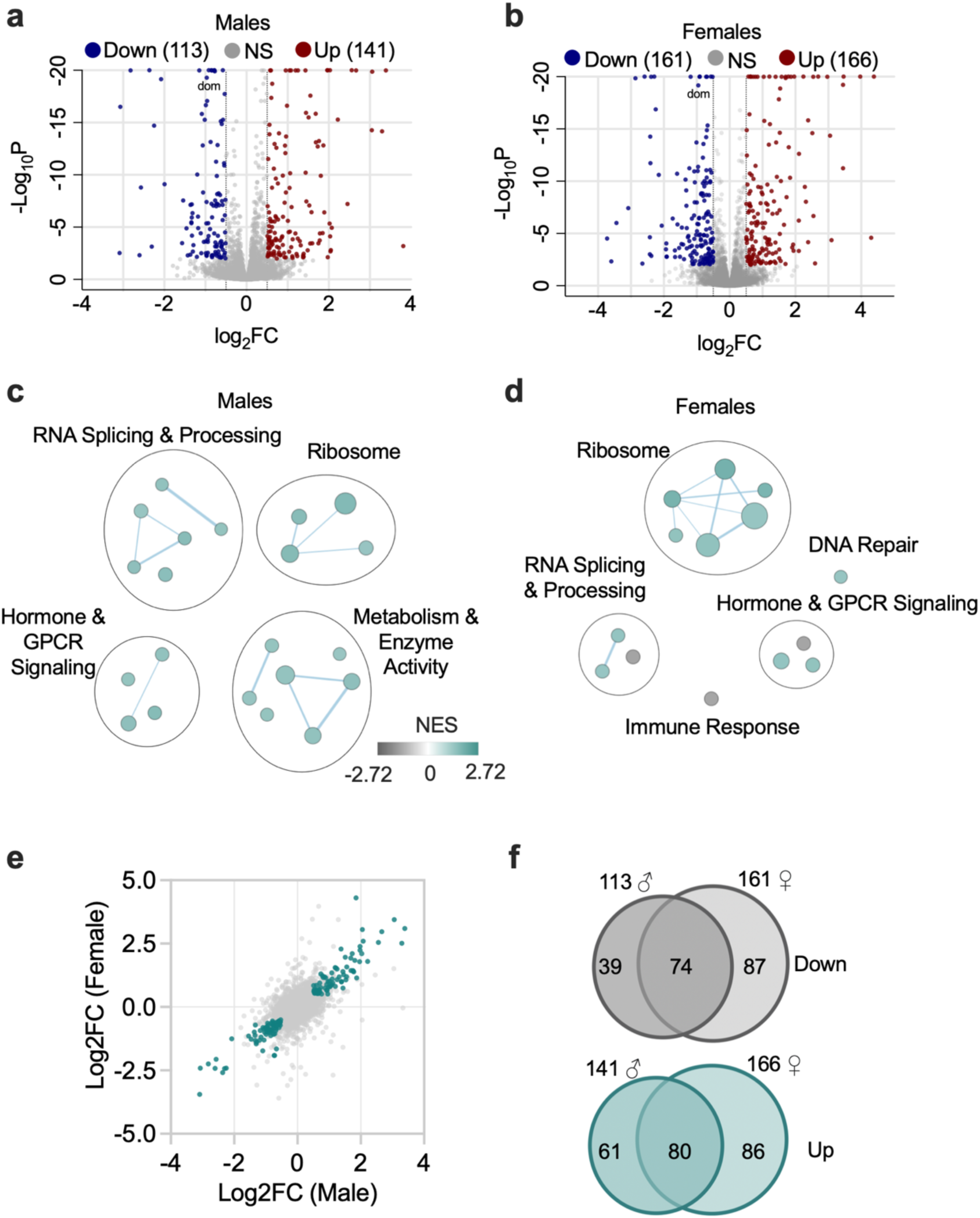
dom^Trojan+/-^ flies exhibit distinct transcriptional profiles in males and females. **a,b**) Volcano plots of DE genes in males (**a**; 113 downregulated and 141 upregulated), and females (**b**; 161 downregulated and 166 upregulated) (n=5 replicates of 5 pooled brains; padj ≤ 0.01 and |log2fold change| ≥ 0.5). Expression of *dom* is significantly reduced in both sexes. **c,d**) GSEA of all DE genes in males (**c**) and females (**d**). **e**) Scatterplot of overlapping significant DE genes in males and females (teal), plotted by log2fold change (overlap significance, Fisher’s Exact Test <0.0001). **f**) Venn diagrams of downregulated (gray) and upregulated (teal) genes in males (left) and females (right). Numbers above circles indicate the total number of DE genes per condition, while numbers within circles denote unique or shared genes across sexes.

**Supplementary Fig. 5:**
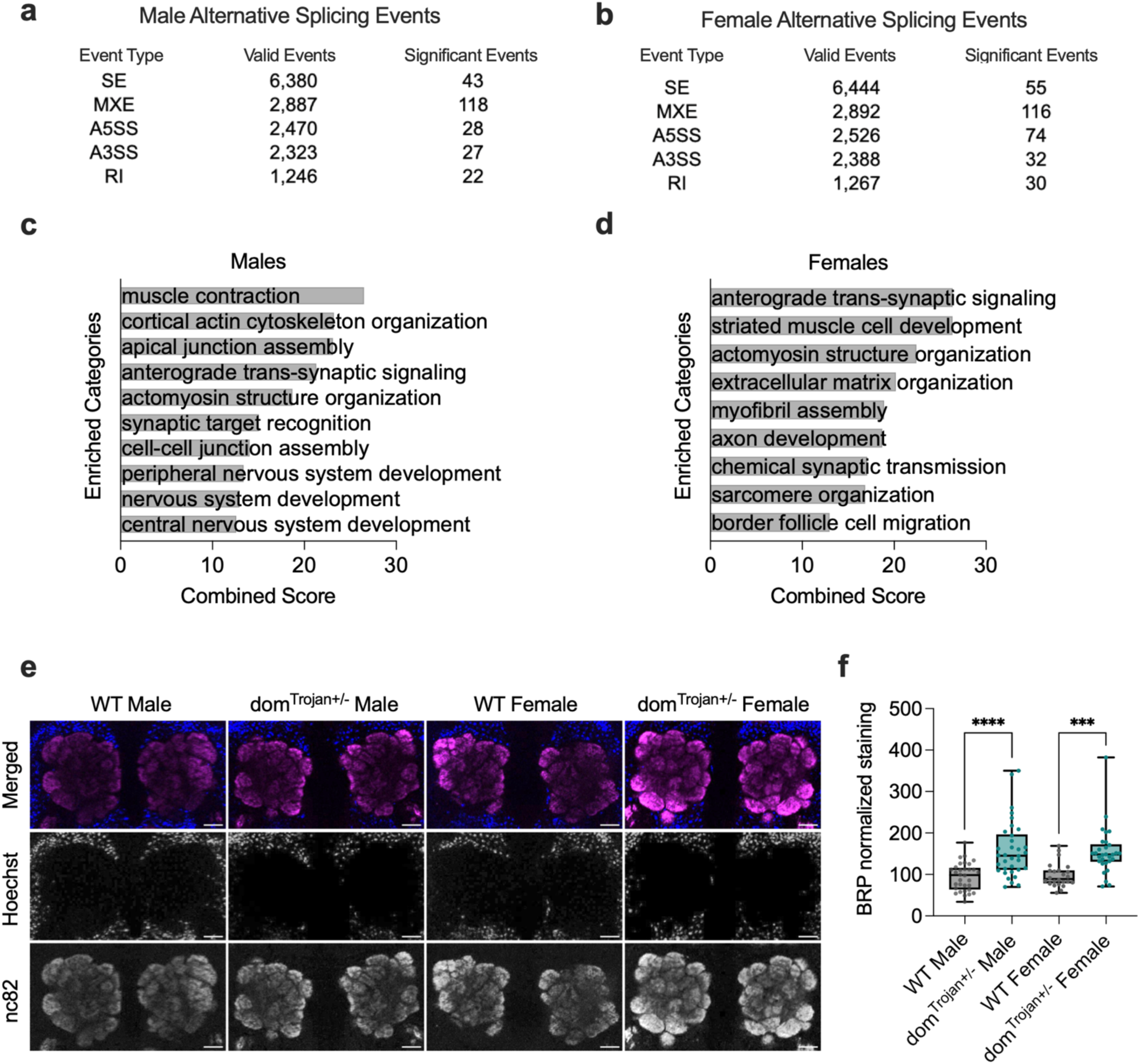
Partial loss of domino alters RNA splicing and synaptic organization in dom^Trojan+/-^ brains. **a,b**)) Alternative splicing events *detected* in male (**a**) and female (**b**) adult dom^Trojan+/-^brains (n=5 replicates of 5 pooled brains), showing total valid events detected by rMATS and significant events (FDR ≤ 0.05 and |IncLevelDifference| ≥ 0.1). **c,d**) ORA of significant splicing events in males (**c**) and females (**d**). **e**) Immunofluorescence images of antennal lobes stained with Hoechst (blue) and nc82 (magenta), with merged images shown for both sexes. Brightness and contrast specifications are consistent across images. Scale bar 20µm. **f**) Fluorescence intensity was measured as mean ROI antennal lobe - mean ROI foramen, with quantification presented as a percentage of the average wild-type intensity (n=23-34; Two-way ANOVA with Šídák multiple comparisons).

## Notes

https://github.com/LempradlLab/drosophila_screen_dom

### Competing Interest Statement

The authors have declared no competing interest.

### Summary of Updates

The screen was updated to completely exclude X-linked genes in the male analysis, with methods revised accordingly. Figures have been updated throughout: Fig. 1 now consolidates all schematics; heatmaps in Fig. 2 and 3 are ranked by significance; Fig. 2 includes PC contribution information; Fig. 3 radar plots feature updated genes; Fig. 4 and Supplemental Fig. 3 provide improved views of individual behavioral parameters; and Fig. 6, 7, and Supplemental Fig. 5 have been re-arranged.

